# Modelling how lamellipodia-driven cells maintain persistent migration and interact with external barriers

**DOI:** 10.1101/2024.09.06.611667

**Authors:** Shubhadeep Sadhukhan, Cristina Martinez-Torres, Samo Penič, Carsten Beta, Aleš Iglič, Nir Gov

## Abstract

Cell motility is fundamental to many biological processes, and cells exhibit a variety of migration patterns. Many motile cell types follow a universal law that connects their speed and persistency, a property that can originate from the intracellular transport of polarity cues due to the global actin retrograde flow. This mechanism was termed the “Universal Coupling between cell Speed and Persistency”(UCSP). Here we implemented a simplified version of the UCSP mechanism in a coarse-grained “minimal-cell” model, which is composed of a three-dimensional vesicle that contains curved active proteins. This model spontaneously forms a lamellipodia-like motile cell shape, which is however sensitive and can depolarize into a non-motile form due to random fluctuations or when interacting with external obstacles. The UCSP implementation introduces long-range inhibition, which stabilizes the motile phenotype. This allows our model to describe the robust polarity observed in cells and explain a large variety of cellular dynamics, such as the relation between cell speed and aspect ratio, cell-barrier scattering, and cellular oscillations in different types of geometric confinements.

**Significance Statement:** Coupling curved membrane proteins to active protrusive forces that arise from recruited actin polymerization, can lead, in the presence of adhesion, to self-organization of a leading-edge cluster and a motile “minimal-cell”. However, this polarized and motile shape can become unstable, and due to fluctuations or interactions with external perturbations transform to an immotile, symmetric shape. Here we couple the spatial organization of the curved active proteins to a global advection of a polarity cue along the cell’s activity axis. Introducing long-range inhibition, the resultant gradient of the polarity-cue stabilizes the motile, polarized “minimal-cell” vesicle. We thereby present a robust model of cell motility that can explain a variety of cellular shape-migration relations, cell-barrier scattering and spontaneous oscillations of confined cells.

## I. INTRODUCTION

During cell migration within the body, such as in development or cancer, cells often have to navigate complex geometries defined by external barriers, tissues and extra-cellular matrix (ECM) fibers [1]. These external constraints and confinements challenge the ability of cells to maintain their internal polarization and exhibit persistent migration. Cellular motility during interaction with complex external constraints and confinement presents an open challenge for our understanding of cell migration.

This process has been explored over recent years using in-vitro experiments where motile cells have been observed while migrating over various topographies and confined within various geometries [2–6]. Theoretical models that describe cellular migration in complex geometries have relied on different coarse-grained cell mechanics approaches [7, 8], phase-field and cellular Potts model (CPM) frameworks [9, 10], or more coarse-grained approaches [11]. These models did not consider the role of curved membrane components and local membrane curvature at the leading edge during lamellipodia-driven cell migration, which is our focus here.

Recently, we have introduced a “minimal-cell” model where a motile phenotype emerges spontaneously due to the coupling between curved membrane complexes (CMC) and protrusive forces that result from the recruitment of actin polymerization to these membrane sites [12, 13]. This coupling can lead to the formation of a lamellipodia-looking protrusion, with a leading-edge cluster of the CMC, and a total resultant force that moves the vesicle in the forward direction (Fig.1A(i)). This simple model has been successful in explaining the origin of several curvotaxis behaviors [14], and since this model is based on only a few physical ingredients, it is very general and applies to many cell types. We note that there are recent experimental indications for the curvature-sensitivity of the leading-edge components of the lamellipodia [15–19], as arises in our model.

**FIG. 1.**
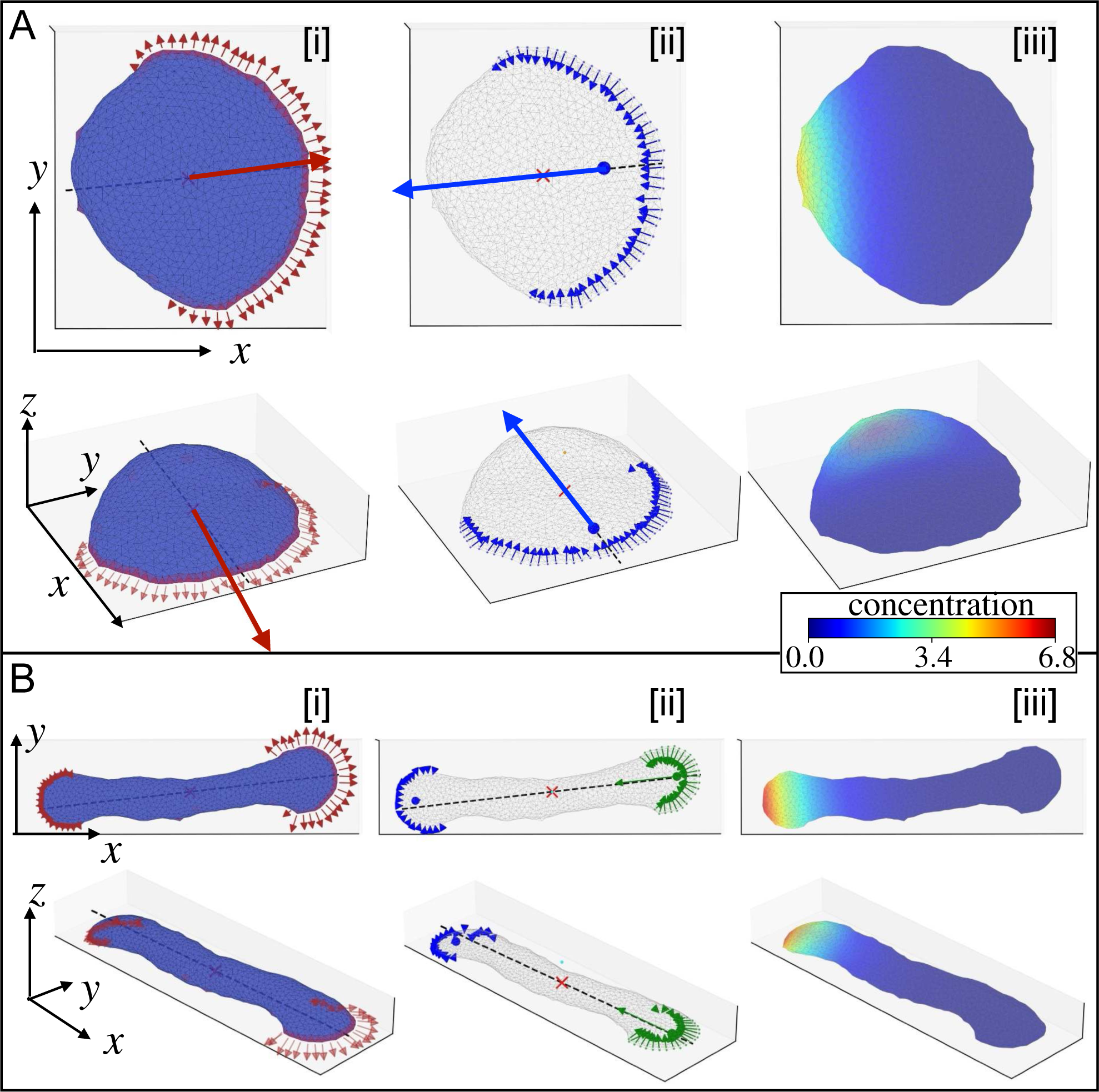
Implementation of the UCSP mechanism and polarity cue advection into the MC simulations of the vesicle shape. This is demonstrated here for two main phenotypes that co-exist in the absence of UCSP [12]: (A) Polar crescent-shaped vesicle 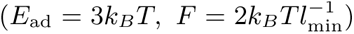: [i] The active force at each CMC site is denoted by a red arrow, directed at the local outwards normal. The big red arrow at the centre of mass (denoted by a red cross) denotes the total active force due to all the active CMC. The black dashed line shows the calculated axis for the net internal flow. [ii] The centre of mass of the protein cluster is denoted by a blue solid circle. The small blue arrows denote the local contribution of each CMC to the global actin retrograde flow (opposite to the direction of the local active force in [i]), and the large arrow denotes the total internal actin-retrograde flow (Eq.1). [iii] The concentration profile of the polarity-cue (inhibitor of the actin polymerization activity), using Eq.(2). (B) Non-polar and immotile two-arc-shaped vesicle 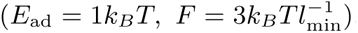.[i]-[iii] as in (A). For each snapshot, we present top and side views.

Nevertheless, within this model the polarized, motile phenotype was rather fragile, and once its leading-edge cluster breaks up, the vesicle irreversibly loses its polarization and forms a non-motile two-arc shape (Fig.1B(i)). Such polarity-loss events can be triggered spontaneously by shape fluctuations or due to the vesicle interacting with external rigid barriers [12]. Similar events of break-up of the leading-edge are observed in motile cells [20–22], however cells have mechanisms that allow them to repolarize and resume their motile phenotype [23, 24].

Here we implement a mechanism for internal cellular polarization [25, 26] into our minimal-cell model, thereby greatly increasing the robustness of the motile phenotype within this model. This allows us to use this model to explain the observed relation between cell speed and shape, as well as the scattering and spontaneous oscillations of motile cells when interacting with external rigid topographical barriers and complex adhesion patterns.

## II. THEORETICAL MODEL

Our minimal-cell model is based on the Monte-Carlo calculation of the dynamics of a closed three-dimensional triangulated self-avoiding vesicle with a spherical topology (Fig.1) [12, 27, 28] (See SI section A for details). Within this model we denote the bare membrane nodes in blue, and the nodes containing the CMC in red (where the active protrusive forces are applied).

The mechanism we implement for the internal polarization of the cell is based on the Universal Coupling between cell Shape and Persistency (UCSP) [11, 25, 26]. Within this model the actin polymerization at the leading edges of the cell gives rise to a net advection of polarity cues across the cell. This advection results in a gradient of the polarity cue along the cell length, and these polarity cue gradients in turn affect the actin polymerization activity at the leading edges, thereby completing a positive feedback that can give rise to spontaneous polarization of the cell. The model was previously mostly implemented in a simple one-dimensional representation of the cell [11, 25, 26], and here we similarly implement it in a simplified manner by projecting the polarity cue gradient and feedback along a one-dimensional axis (ignoring hydrodynamic flow patterns - see in a two-dimensional cell [29]). We further note that the front-back inhibition may arise in cells due to other types of long-range inhibition, and one can view our implementation as an example of a large class of intra-cellular mechanisms that stabilize cell polarization [30–35].

First, we calculate the axis of the net retrograde flow within the vesicle, for the instantaneous configuration of membrane shape and CMC clusters. This is shown in Fig.1, and more details are given in SI section B.

The magnitude of the net retrograde flow is related to the active cytoskeleton force exerted by the CMC (Fig.1(A,B)[i]) by a positive coupling factor *β*. Each CMC contributes to the total actin retrograde flow, which is oppositely directed to the active protrusive force

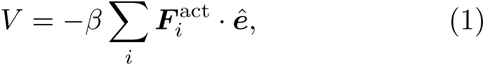

where, ***ê*** is the direction of net retrograde flow along the calculated axis in such a way as to have a positive *V* (Fig.1(A, B)[ii]), and 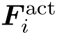 is the active actin-driven protrusive force due to a CMC at the *i*th vertex, pointing at the outwards normal.

Using the calculated retrograde flow, we calculate the polarity cue distribution along this axis, given by,

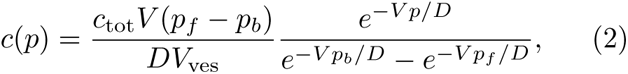

as shown in (Fig.1(A,B) [iii]). Here, *p* is the projection of a vertex position with respect to the centre of mass on the axis of retrograde flow, *V*_ves_ is the volume of the vesicle, and *D* is the diffusion coefficient of the inhibitory polarity cues. *p_f_* and *p_b_* are the projections of the vertices at the front and the back of the cell with respect to the retrograde flow axis. The quantity *c_tot_* denotes the total amount of the polarity cue: 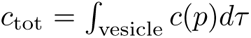. In our model the unit of the actin retrograde flow *V* is given by *D/l*_min_, since its only the ratio *V/D* that appears in Eq.2. The force has units of 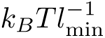, and therefore the coupling parameter *β* has units of 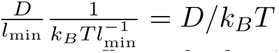.

The polarity cue distribution in turn affects the local strength of the protrusive force induced by actin polymerization at the location of each CMC, according to the following steady-state of first-order kinetics

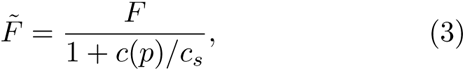

where *c_s_*is the saturation concentration for this inhibitory reaction. This feedback means that in the region with the higher concentration of inhibitor the CMC induce weaker protrusive forces, and contribute less to the total actin retrograde flow (Eq.1). This is clearly demonstrated in Fig.1B.

The modified actin polymerization forces (Eq.3), and the modified contribution to the actin retrograde flow (Eq.1), are then used to calculate again the polarity cue profile (Eq.2) until these repeated iterations converge to within some fixed error threshold of 0.1%. The MC simulation is then allowed to proceed with the modified CMC forces, until the vesicle shape has changed by more than some threshold value 10%, and the UCSP calculation is updated, whereby a new axis for the net flow is calculated and the whole process is repeated.

The examples shown in Fig.1 demonstrate the principles of this procedure for two typical shapes that form spontaneously in our model: the motile crescent shape and the non-motile two-arc shape. Both shapes co-exist in the same parameter regime, when we do not implement the UCSP calculation. In both cases, we indicate the identification of the axis of the net retrograde flow, and the resulting concentration profile of the polarity cue. We show here the first iteration of the UCSP calculation.

In the UCSP model we found that spontaneous polarization of the cell occurs when the coupling parameter between the actin flow and the asymmetry in the polarity cue (*β*) is larger than a critical value [25], which also depends on the cell length [26]. We show in the SI a similar transition from a non-motile to a motile (polarized) state above a critical value of *β* (Fig. S-1).

All the vesicles used in this work are composed of *N* = 1447 vertices to create the triangulated membrane. We used the time unit as 20000 Monte Carlo steps and the protein percentage is *ρ* = 3.45% if not specified (For pancake shape, we used *ρ* = 5.53% in Fig.3). We used the following UCSP parameters throughout the paper: *c*_tot_ = 4000, *D* = 4000, and *c_s_* = 1, and the CMC binding energy was fixed at *w* = 1 *k_B_T*.

## III. RESULTS

We now demonstrate the results of implementing the internal polarization mechanism (UCSP) on the motility dynamics of our minimal-cell model.

### A. UCSP mechanism stabilizing the motile phenotype

We have previously shown that the crescent shape is transiently stable and can make a spontaneous transition to a two-arc shape, since both shapes coexist in the same parameter regime [12]. This instability occurs faster for large protrusive force *F* and weak adhesion *E*_ad_, and is shown in Fig.2. In Fig.2A we demonstrate that in the absence of UCSP (*β* = 0 *D/k_B_T*), the leading-edge cluster can spontaneously break into two, whereby the vesicle changes to the two-arc shape and motility is irreversibly lost. As the coupling between the asymmetry in the polarity cue and the actin flow (*β*) increases, the UCSP mechanism can stabilize the crescent-shaped vesicle and suppress the transition to the non-motile two-arc shape. As *β* increases the polarity cue profile becomes sharper from back to front (Fig.2B), and the net retrograde flow remains more stable (Fig. S-2), preventing the transition to the two-arc shape.

**FIG. 2.**
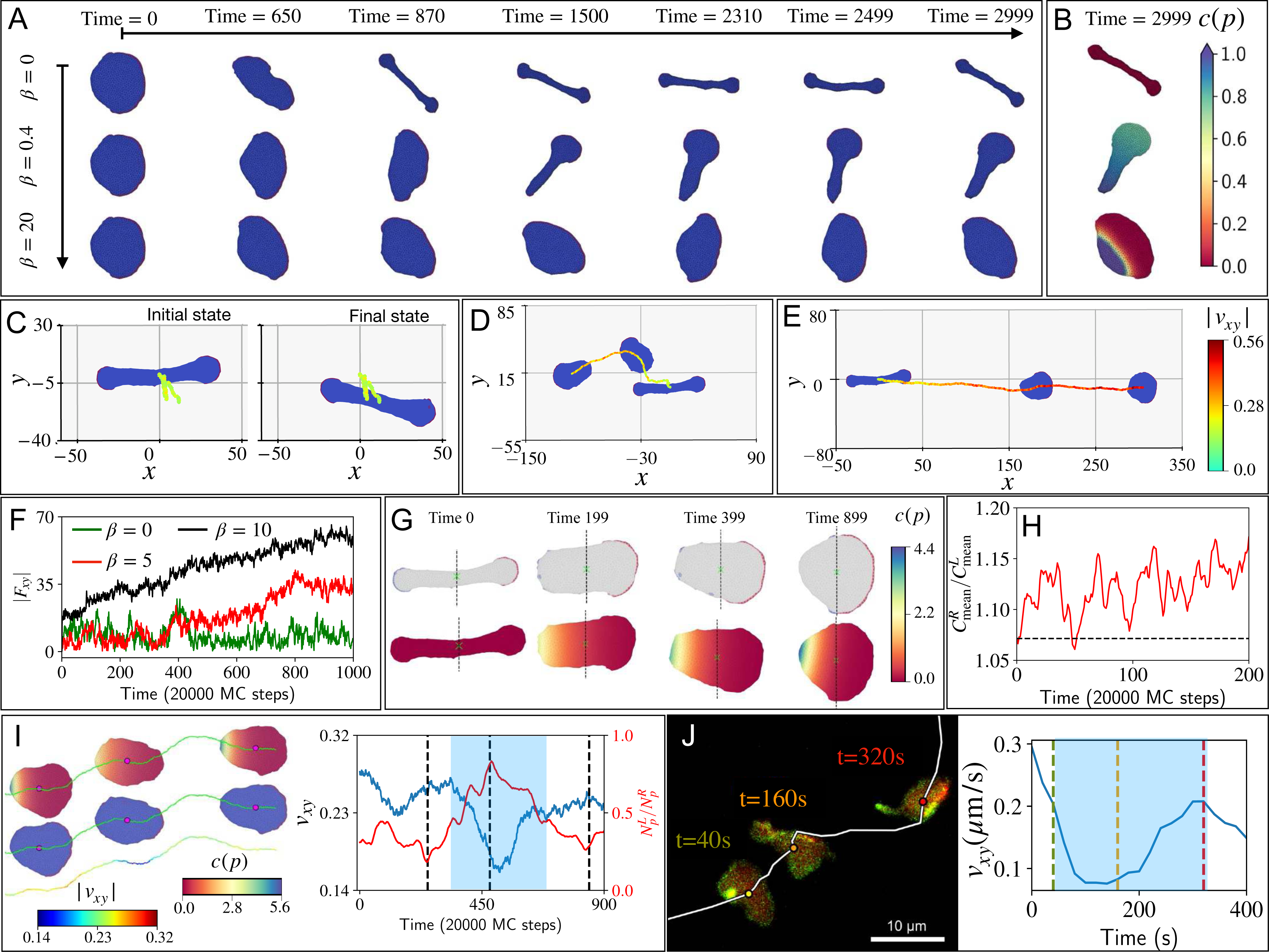
Stabilization of the crescent-shaped, polar vesicle by the UCSP mechanism. (A) Examples of the dynamics of the motile vesicle for different values of the UCSP coupling strength *β*. In the absence of UCSP (*β* = 0 *D/kBT*) the crescent shape is transiently stable, and thermal fluctuations break it into the two-arc shape [12]. As the coupling strength *β* increases, the instability is delayed or allows the polar state to be partially recovered (*β* = 0.4 *D/kBT*). When the coupling is strong (*β* = 20 *D/kBT*) the crescent shape becomes absolutely stable. In order to maintain a constant maximal active force magnitude of 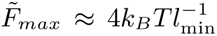, we used *F* = 4, 7.1, 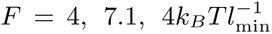 for the cases of *β* = 0, 0.4, 20.0 *D/kBT* respectively (the adhesion parameter *E*_ad_ = 1 *k_B_T*). See Movie S-1. (B) The concentration profile of the polarity cue along the UCSP axis at the final times of the cases shown in (A). (C-E) The transition of a two-arc shape to a crescent polar-shaped vesicle (See Movie S-4). The trajectories indicate the speed of the vesicle using the heatmap. We started the simulations with a two-arc-shaped vesicle using 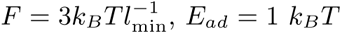. (C) In the absence of UCSP coupling (*β* = 0 *D/k_B_T*), the two-arc shape is stable. (D,E) For higher coupling strength (*β* = 5, 10 *D/kBT*), the UCSP mechanism breaks the symmetry of the two-arc shape and makes a transition to the crescent, motile shape (shown at times 0, 699, 999). (F) The planar force magnitude *Fxy* is shown for *β* = 0, 5, and 10 in units of *D/kBT* with green, red and black respectively. (G) The snapshots of the shape transition from a two-arc to a crescent shape for *β* = 10 *D/kBT* (the case *β* = 10 *D/kBT* shown in E). The CMC on the right (*xi > x*CM) and left (*xi < x*CM) of the centre of mass are marked by red and blue respectively. The concentration profile of the inhibitory polarity cues is shown. (H) The time evolution of the ratio of the average curvature of the CMC on the right and left of the centre of mass. (I) Increased amplitude of thermal fluctuations by decreasing the bending rigidity *κ* = 15*kBT*, giving rise to spontaneous transition between polar and non-polar shapes. We set the coupling parameter *β* = 20 *D/kBT*. The polar-shaped vesicle makes a transition to a nearly two-arc shape and back to a motile crescent shape (See Movie S-3). The two-arc shape corresponds to the dip in the planar speed plot (blue), and a peak in the ratio of left/right number of CMC with respect to the c.o.m. along the polarity axis (red). (J) Experimental data is showing a *D. discoideum*cell undergoing a transition between polar and non-polar (cells scaled down by a factor of 2, labelled with LifeAct-GFP and PHcrac-RFP). The snapshots are shown at 40s, 160s and 320s (See Movie S-5). The planar speed *vxy* shows a similar dip when it becomes a two-arc. The dashed lines correspond to the time at which we showed the shape of the cell from the experiment.

Next, we study the UCSP-induced polarization of the two-arc shape (Fig. 2). We start the simulation with a non-motile two-arc-shaped vesicle, which is stable in the absence of the UCSP mechanism (Fig. 2C). We then switch on the UCSP mechanism and demonstrate how the vesicle becomes more polarized, crescent-like and motile as *β* increases (Fig. 2D-E). The total active force increases as the cell polarizes and transforms from the two-arc to the crescent shape, as shown in Fig. 2F.

The polarization process proceeds as follows: the polarity cue concentration peaks at one end of the two-arc shape (Fig. 2G), inhibiting the protrusive forces in this “losing” CMC cluster. This reduction in the amplitude of the protrusive forces causes this cluster to lose its high curvature (Fig. 2H), which then loses its stability, breaks up and its CMC diffuse to join the ”winning” cluster at the opposite end of the two-arc shape (Fig. 2G), where the large protrusive force maintains a high curvature at the leading edge. This process thereby converts the two-arc shape into the crescent motile phenotype with a single large CMC cluster (Fig. 2D,G). Clearly, there is a transition regime of values *β ∼* 1 *−* 5 in units of *D/k_B_T* above which the minimal-cell can robustly repolarize following its transition to the non-motile two-arc shape.

In order to observe more dynamic transitions in polarity, which are often observed in living cells, we need to allow for larger fluctuations in our system. By reducing the bending rigidity parameter from *κ* = 20 *k_B_T* to *κ* = 15 *k_B_T* (and using an active force parameter 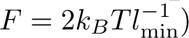 we induce larger amplitude shape fluctuations, effectively emulating the large level of metabolic noise observed in cells. Now, we observe spontaneous shape changes (See Movie S-3) from crescent-shaped polar to non-polar nearly two-arc-shaped vesicle, as shown in Fig. 2I. The planar speed *v_xy_*dips when the vesicle transitions to a two-arc shape (blue line in Fig. 2I). To quantify the distribution of the CMC clusters we plot the ratio 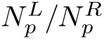 of the number of proteins on the left and right sides of the centre of mass, along the polarity axis (red line in Fig. 2I). As this ratio approaches 0 (1) it implies a more (less) polar vesicle, and indeed this ratio 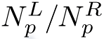 reaches its maximum when the planar speed is lowest. This sequence of polarity loss, and recovery, is observed in experiments with *D. discoideum* cells, where spontaneous switches between a crescent-shaped polar mode of locomotion and a non-polar state were reported [36] for an example see Fig. 2J (See Movie S-5). A similar dip in planar speed *v_xy_* is observed experimentally when the cell takes a two-arc shape. Note that in the absence of UCSP (*β* = 0 *D/k_B_T*), the larger thermal fluctuations break up the motile vesicle, which quickly becomes two-arc (see Movie S-2).

We further investigated the interplay between cell shape and polarization by starting with a non-polar vesicle where the CMC form a circular leading edge around the entire shape (using a higher CMC concentration, Fig.3A). In the absence of UCSP this “pancake” shape is stable and non-motile [12]. We then apply a transient external force field, which emulates the effect of blowing fluid at high pressure on the vesicle (see SI section E for the details of this external force field), resulting in the deformation of the pancake into a crescent shape (Fig.3B). This simulation is motivated by the experiments that demonstrated the conversion of a non-motile circular cell fragment into a motile phenotype by deforming it into a crescent shape by an applied shear flow [37].

**FIG. 3.**
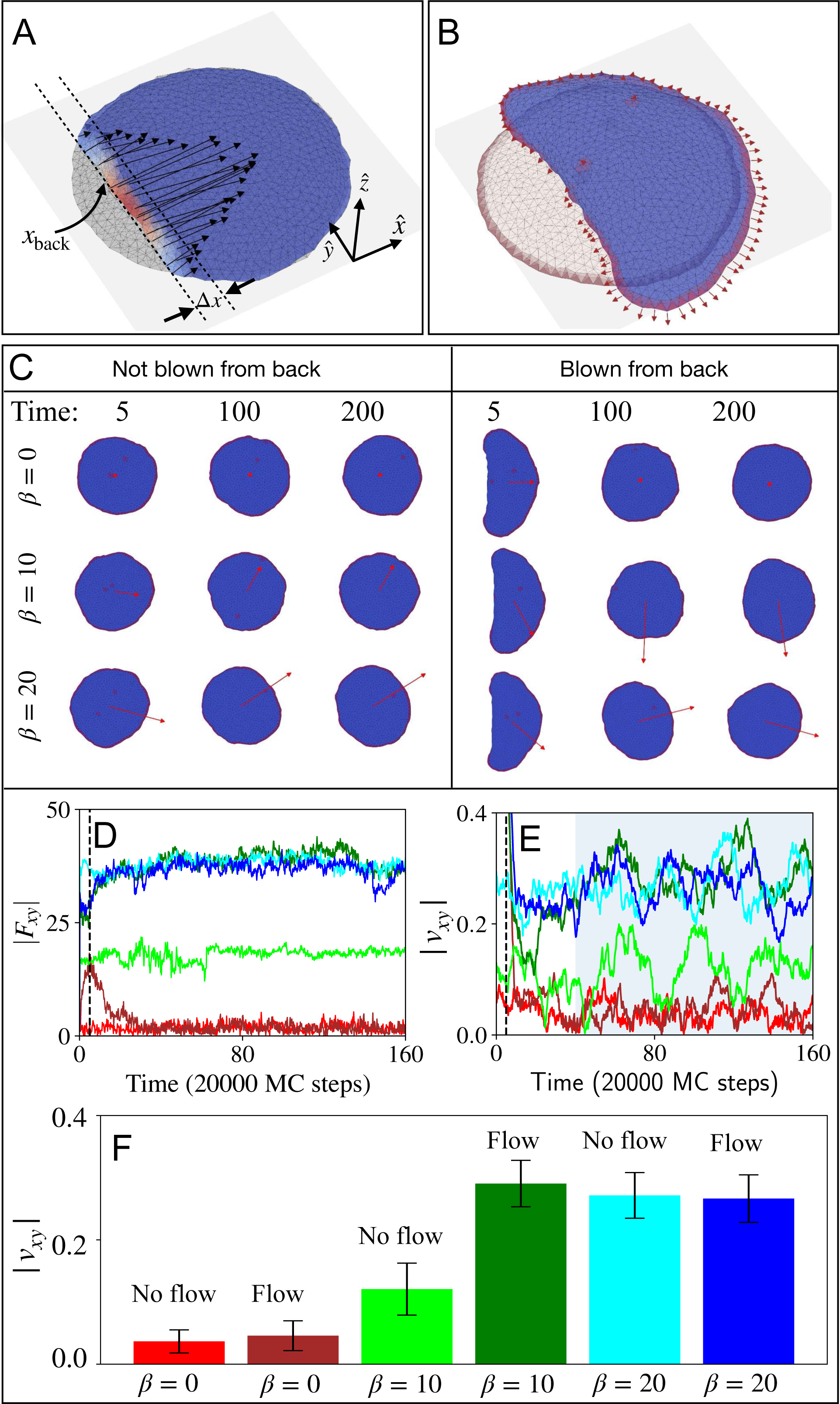
The relation between the vesicle’s initial shape and polarization. (A,B) The initial state of a spread vesicle, with a circular leading-edge cluster (grey background image). An external force field (black arrows in (A)) is applied to deform the vesicle, mimicking the effect of a strong fluid flow in the *x̂* direction. The force is acting within a range of Δ*x* of the vesicle’s rear. The strength of the force has a Gaussian form (heatmap). (B) The final deformed, crescent shape when the external force field is turned off. The leading-edge cluster is broken-up in the region deformed by the external force, due to the membrane losing its high curvature. The local active forces are indicated by the red arrows. (C) Vesicle shapes at different times for different coupling strengths (*β* = 0, 10, 20 in units of *D/kBT*), for the pancake and the deformed crescent shapes (B). (D,E) The time dependence of the total planar active force 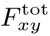 and velocity *vxy*. The colour code of the lines is elaborated in (F), where we showed a bar plot of *vxy* averaged over a time window indicated by the shaded region in (E). The parameters used: parameter protein density *ρ* = 5.53%, adhesion strength *E*_ad_ = 3*k_B_T* and active force parameter *F* = 2, 2.5, and 2 in units of 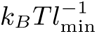 for the cases *β* = 0, 10, and 20 in units of *D/kBT* respectively to maintain the maximum force at one vertex 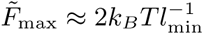.

In Fig.3(C-F) we demonstrate the effect of such a transient shape deformation on the polarization of the vesicle, for different values of the UCSP coupling parameter *β*. When the transient external force field is switched off, we implement the UCSP mechanism, and follow the consequent evolution of the vesicle’s polarization. We find that the shape deformation leads to a small polarization for small (or no) UCSP coupling strength, which decays over time. Initially the crescent shape disrupts the CMC cluster in the region facing the external force field (Fig.3B), as its curvature becomes small or negative (concave). However, this polarization of the CMC and the total force is short-lived and the vesicle resumes the pancake shape, with decaying polarization.

At large values of *β* the UCSP coupling is strong enough to break the symmetry and polarize even the pancake shape without the transient crescent shape deformation. There is however an intermediate regime (for example *β* = 10 *D/k_B_T*), where we find that the transient crescent shape deformation enables the system to attain a persistent high polarity form, which it can not reach spontaneously from the pancake shape. This observation demonstrates that within our model there can be coexistence of long-lived low and high polarity phenotypes which depend on their initial shape, as observed in experiments [37].

The correlation between cell polarization, as manifested by the cell velocity, and cell shape were experimentally measured for different types of motile cells [38, 39]. We use our model to explore this relation by simulating vesicles with different densities of CMC and strength of the UCSP coupling (Fig.4). For each combination of CMC concentration and *β* we plot in Fig.4A the instantaneous speed and aspect ratio of the vesicle’s projection on the *x − y* plane (calculated with respect to the direction of motion). In Fig.4(B-E) we show typical snap-shots of the vesicle shape and CMC cluster along the vesicle leading edge. Remarkably, the relation that we obtain between speed and aspect ratio (Fig.4A) exhibits the same step-like behaviour observed in experiments, including the large scatter [38, 39].

**FIG. 4.**
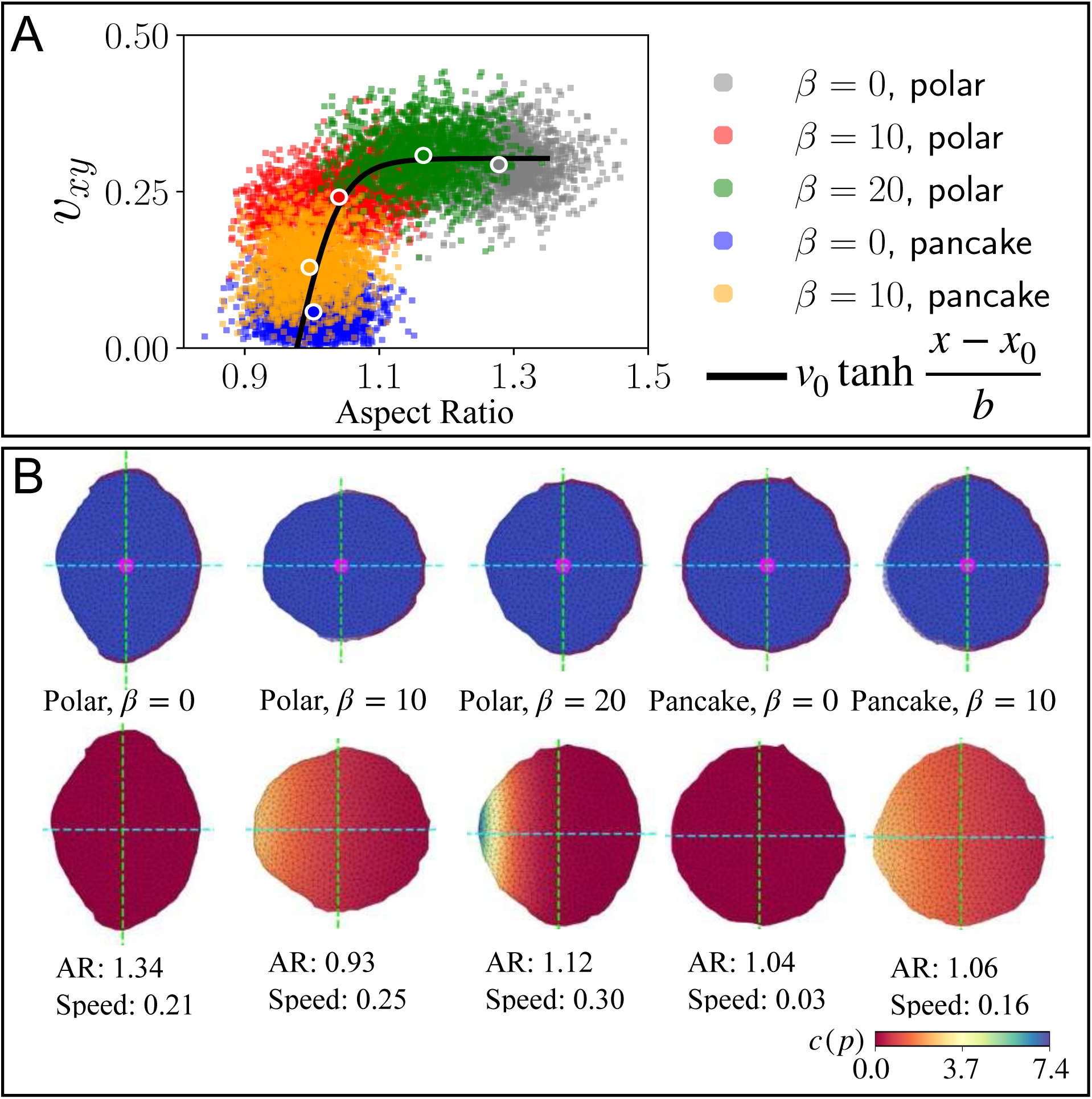
Relation between the vesicle’s aspect ratio and speed. (A) Scatter plot of the instantaneous speed and aspect ratio of the vesicle for four different cases: (i) polar vesicle, *β* = 0 *D/kBT*, (ii) polar vesicle, *β* = 10 *D/kBT*, (iii) polar vesicle, *β* = 20 *D/kBT*, (iv) pancake-shaped vesicle, *β* = 0 *D/kBT* in grey, red, green, and blue respectively. The black line shows the fitted-curve with a saturating functional form *v*0 tanh *b*(*x − x*0) through the average coordinates of each cluster. We find *v*0 = 0.3 *±* 0.0014*, b* = 0.06 *±* 0.002, and *x*0 = 0.977 *±* 0.0003 using python scipy package. (B) Typical shapes of the vesicles for the different cases, and the polarity cue distribution profiles. The CMC density is 3.45% and 5.53% for the polar and the pancake shapes respectively.

As expected, the vesicles with the lower concentration of CMC maintain a more polar form, with and without the UCSP. In fact, the vesicles with UCSP of intermediate coupling strength (*β* = 10 *D/k_B_T*, red dots) have a lower aspect ratio compared to no UCSP (*β* = 0 *D/k_B_T*, grey dots) since the forces exerted sideways, which stretch the vesicle perpendicular to its direction of motion, are inhibited in this case. With larger aspect-ratio a larger proportion of the forces are oriented in the direction of motion, allowing for higher average speed. When the CMC form a circular cluster, the cell can still polarize for significant coupling strength (*β* = 10 *D/k_B_T*, similar to Fig.3), but with lower speeds due to significant forces acting opposite to the direction of motion.

Our model therefore allows to explain the aspect-ratio-speed relation, through the processes by which the active forces at the leading edge both propel and deform the cell, with the efficiency of the propulsion (speed) dependent on the shape and the resulting distribution of the leadingedge cluster around the cell. Note that our model at present does not include the contractile forces that occur at the rear of polarized cells and cell fragments [37–40]. Such contractile forces, localized at the cell rear, can deform polarized cells and cell fragments into the crescent shapes observed in experiments.

### B. Interactions with barriers and confinements

We now explore the dynamics of our minimal-cell model, incorporating the UCSP, when interacting with external barriers and confinements. In the absence of the UCSP (*β* = 0 *D/k_B_T*), we found that when our motile vesicle impinges on a rigid wall barrier, it loses its polarity and converted to the non-polar two-arc shape [12] (Fig.5A). This happens in our model due to the membrane flattening against the rigid barrier, losing its high curvature along the leading-edge and the subsequent migration of the highly curved CMC to the nearest free membrane on either side along the barrier (See Movie S-6). While real cells do transiently lose or diminish their polarity when scattering off barriers, they can recover their motility and migrate away [23, 24]. Similar behaviour is observed when cells collide [41], giving rise to cell-cell scattering and a form of contact inhibition of locomotion (CIL) [42].

**FIG. 5.**
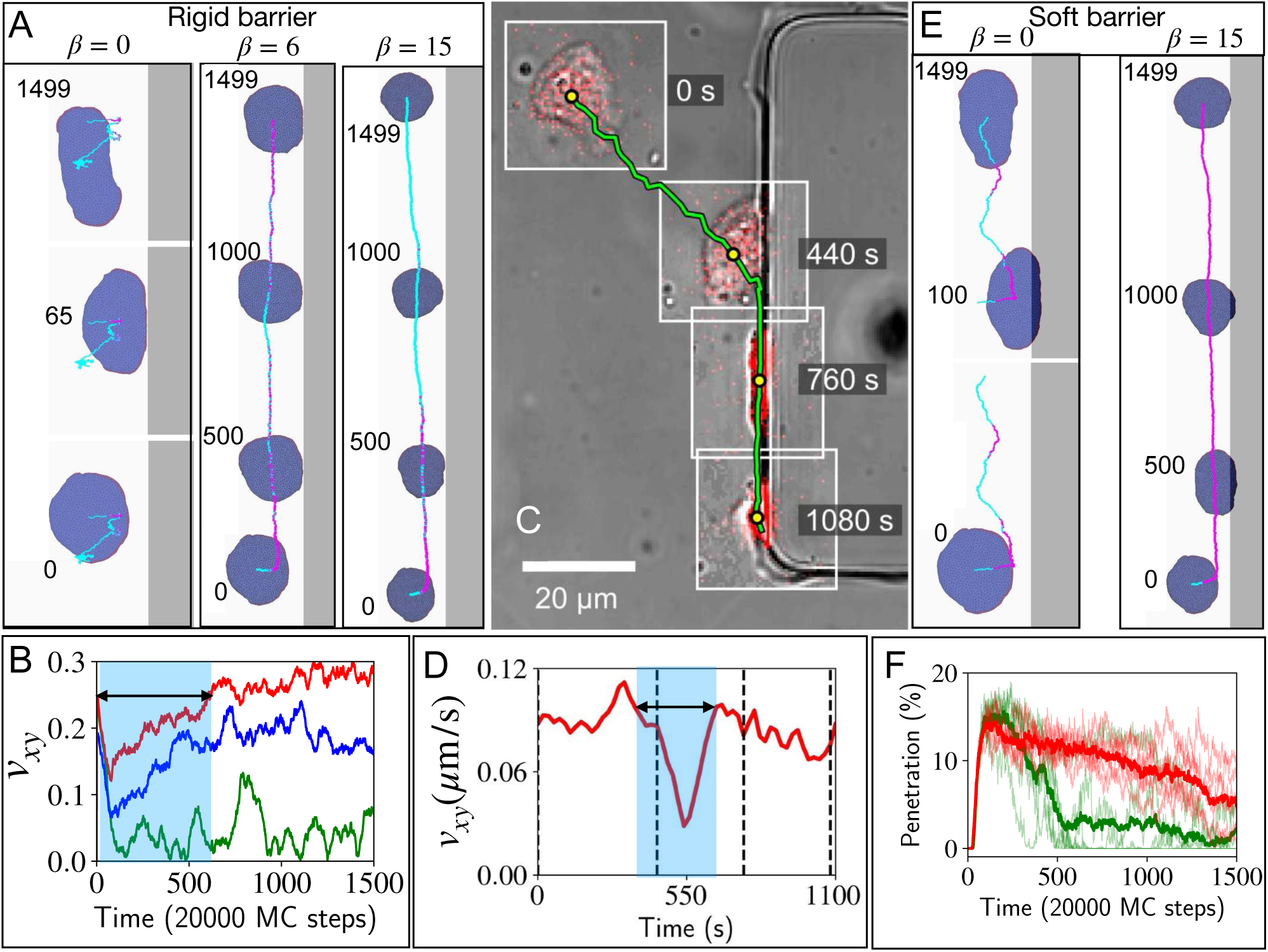
Scattering and repolarization of a cell hitting a rigid barrier. (A) The trajectory and shapes of the vesicle hitting a rigid barrier (grey region) for three different values of the coupling parameter *β* = 0, 6, 15 in units of *D/kBT* (See Movies S-6, S-7). (B) The planar speed *vxy* from the simulation is shown for the rigid barrier. We used green, blue and red colours for *β* = 0, 6, 15 in units of *D/kBT* respectively. (C) Trajectory and shapes of a *D. discoideum* cell migrating and hitting a rigid barrier in the experiment (see Movie S-9). The snapshots from different times (insets with a white border) where overlaid on the bright field channel, to visualize the barrier. Only the red channel (PHcrac) and bright field channel are shown. (D) The planar speed *vxy* of the *D. discoideum* cell from the experiment. (E) The trajectory and shapes of the vesicle hitting a soft barrier (spring constant 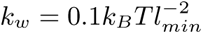, grey region) for two values of the coupling parameter *β* = 0, 15 in units of *D/kBT*. (F) Penetration percentage is plotted over MC steps in the case of soft barriers for *β* = 0, 15 *D/kBT* in green and red respectively. The trajectories in (A) and (E) are coloured cyan if the vesicle is not in contact with the wall, or magenta if it is touching (within *s* = 0.15 *lmin* of the rigid wall) or penetrating the soft wall region. We used 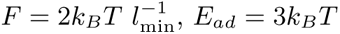.

In Fig.5A we demonstrate the trajectory of the same vesicle in the presence of UCSP (*β* = 6, 15 *D/k_B_T*) (See Movie S-7 for *β* = 15 *D/k_B_T*). While the speed and polarization transiently diminish when the vesicle hits the barrier (Fig.5B, Fig. S-5A), it recovers and the vesicle migrates away. In Fig. S-6 we give more examples of cell-barrier scattering simulations at various angles.

In Fig.5C, we demonstrate an experimental trajectory of a *D. discoideum* cell that hits a rigid barrier, and continues to slide along it. The speed of the cell is shown in Fig.5D. During the interaction with the barrier, the cell transiently loses its motility (blue shaded region) and then recovers the speed again as it slides along the barrier edge. This is very similar to the behavior in our simulations (Fig.5A,B).

Recently it was observed that when the cells can partially penetrate the barrier, they often get trapped at the barrier for a significant period of time, before escaping away [43]. We simulate the effect of a soft barrier by allowing the vesicle to move into the barrier (Fig.5E, F), which exerts a spring-like restoring force on each membrane node, proportional to the penetration distance (see SI section F for more details).

Comparing to the hard-wall case, we find that even without the UCSP mechanism (*β* = 0 *D/k_B_T*) the motile vesicle maintains its polarity when hitting the soft wall (Fig.5E). This is facilitated by the membrane maintaining its high curvature along its leading-edge, thereby preventing the CMC cluster from breaking up into the two-arc configuration (See Movie S-11). After spending some time stuck against the wall, the vesicle spontaneously rotates and migrates away (with significantly diminished polarization, Fig. S-5B, C). In the presence of UCSP, the cells get stuck penetrating the barrier for longer times as the coupling strength increases (Fig.5F), before migrating away (See Movie S-12).

A more extreme scattering configuration is presented in Fig.6. Motivated by experiments we let our motile vesicle hit the triangular tip of a square-shaped barrier, edge on (the shape of the triangular tip is explained in SI section G.1). As expected, in the absence of UCSP, the vesicle loses its polarity to the immotile (See Movie S-13) two-arc shape (Fig.6 A). As the values of *β* increase the UCSP mechanism which allows the vesicle to recover its polarized shape following the scattering with the barrier, and continue its migration (See Movie S-14). In Fig.6D we show similar scattering events in experiments using *D. discoideum* cells hitting PDMS barriers(See Movie S-15). As in the model, the cells can lose their polarity, form two competing protrusions with very large shape elongation (similar to the two-arc shape), and recover their polarity.

**FIG. 6.**
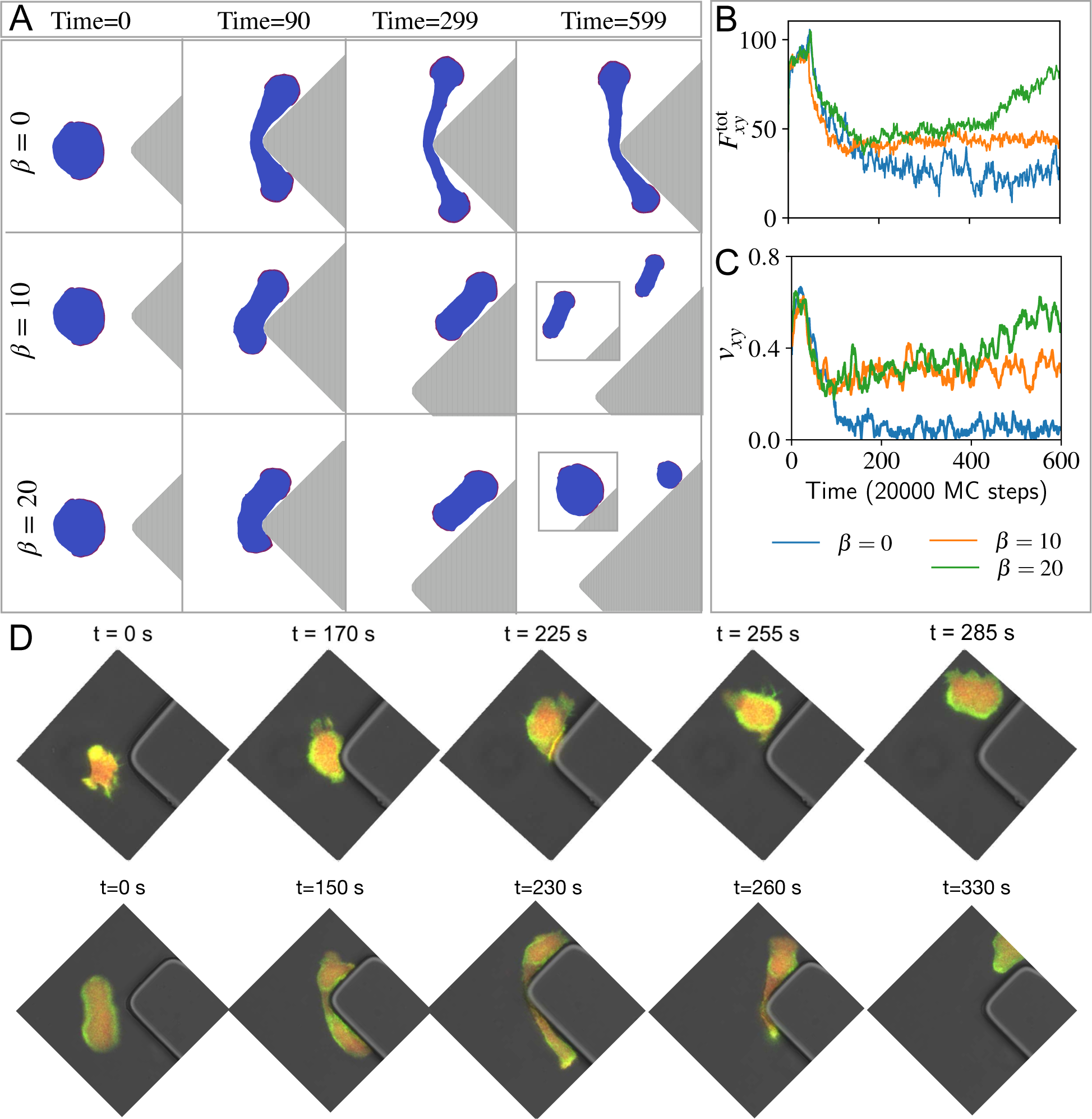
Scattering of a vesicle and a cell from a triangular shape. (A) Snapshots of the vesicle, initially in the polar state, when hitting head-on a rigid boundary with a triangular shape, for four different coupling strengths *β* = 0, 10, 20 *D/kBT*. We used the active force 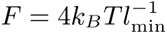 and adhesion strength *E*_ad_ = 3*k_B_T*. For *β* = 0 *D/k_B_T*, we can see that the vesicle loses its polarity. For intermediate coupling, *β* = 10 *D/kBT*, the repolarization of the vesicle is partial, while it is complete for strong coupling, *β* = 20 *D/kBT*. (B,C) The total planar active force 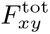 and velocity *vxy*, respectively. This demonstrate the loss of polarization, and its recovery as function of time for the different values of *β*. (D) Snapshots from experiments showing similar dynamics of motile *D. discoideum* cells scattering off the triangular tip of a PDMS barrier. The cropped regions of interest (ROIs) have been rotated to display the same orientation as the examples shown in (A). The ROI in the first and second row is 50*×* 50 *µ*m and 75*×* 75 *µ*m respectively.

The most elaborate test of our vesicle’s motility under confinement is shown in Fig.7A. Here, we consider a dumbbell-shaped region surrounded by rigid barriers. The configuration of this confinement is motivated by experiments that have demonstrated spontaneous cellular oscillations within this system [44, 45]. In these experiments it was found that cells spontaneously oscillate along the dumbbell pattern, and our model displays very similar behaviour (Fig.7A,B). Within our model, the origin of this oscillatory behaviour is explained in Fig.7C: As the vesicle moves to the right, its leading edge reaches the rigid walls of the confining barrier, where the leading edge loses its high curvature (Fig.7D), and the leading edge cluster breaks into two (or more) clusters on either side, protruding mainly along the perpendicular directions. The contributions of these clusters to the global retrograde actin flow approximately cancel each other, and the global flow along the long axis of the pattern (*x*-axis) becomes dominated by the trailing edge CMC cluster and switches direction. As the flow direction switches, so does the polarity cue gradient and the activity of the CMC becomes strong in the new leading edge and weak at the new trailing edge, and the cell moves towards the opposite end of the dumbbell, and so on (See Movie S-16).

**FIG. 7.**
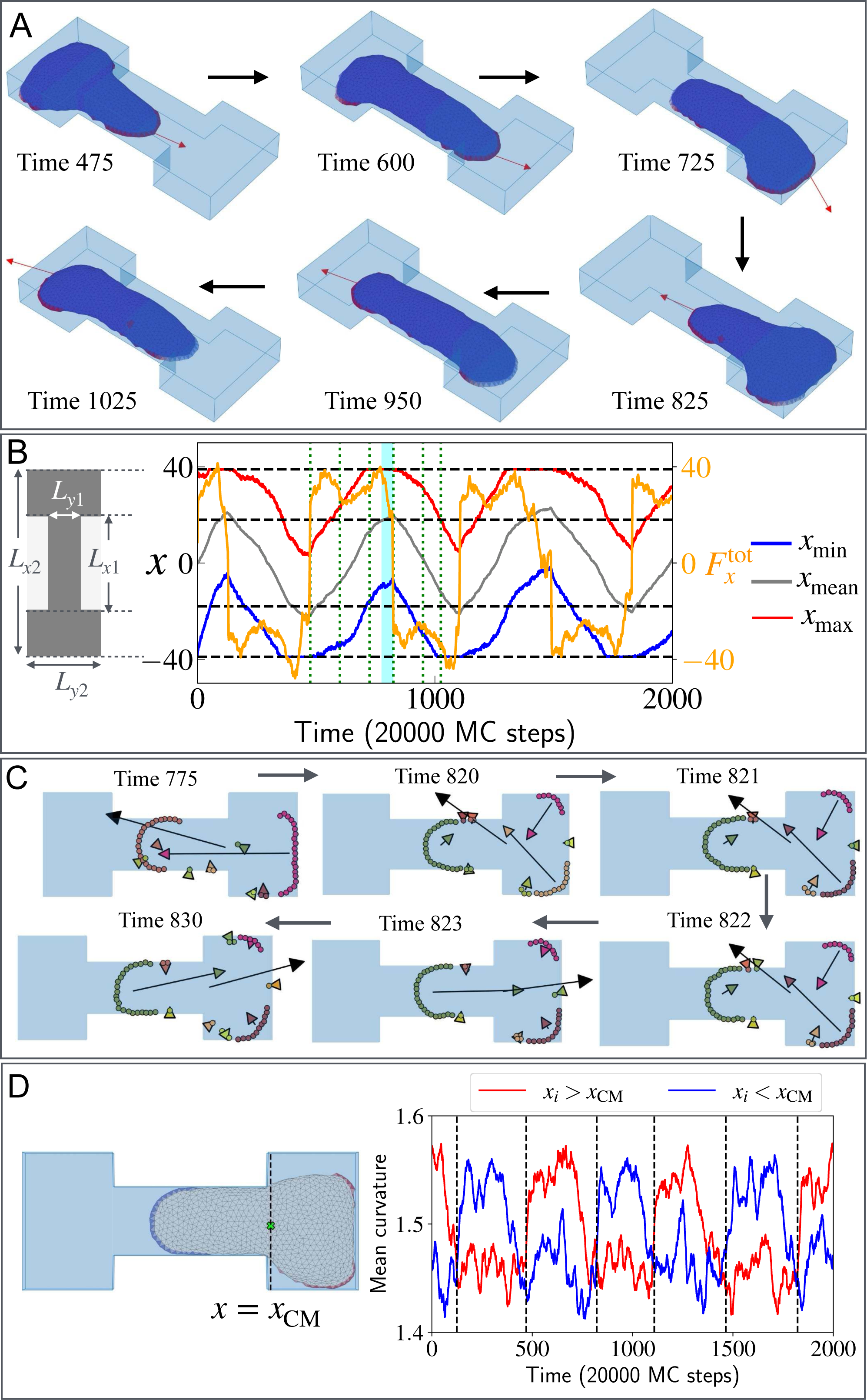
Oscillations within a dumbbell topographic confinement. (A) Snapshots of the vesicle within the dumbbell-shaped confinement at time 475, 600, 725, 825, 950, and 1025 in the units of 20000 MC Steps during a complete oscillation between the two chambers. The total active force 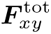 (excluding the *z* direction) is shown with the red arrow for each snapshot. (B) The oscillation along the *x* coordinate of the vesicle over time. The *x*min, *x*mean, and *x*max of all the vertices of the vesicle are shown in blue, grey, and red solid lines respectively. Black dashed lines indicate the dimensions of the dumbbell shaped-confinement, where, *Lx*1 = 36, *Lx*2 = 78, *Ly*1 = 16, and *Ly*2 = 32 in the units of *l*min. The time evolution of the *x* component of the total force 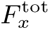 is shown in orange (scale on the right). Green vertical dotted lines denote the times of the snapshots in (A). (C) Different protein clusters are shown in different colours on a *x − y* plane from the top view during the polarity flip. We indicate the actin flow contribution from each cluster in the corresponding coloured arrow at the mean position of that cluster. The total actin flow is indicated using a black arrow at the centre of mass of the vesicle. We use the adhesion strength *E_ad_* = 3*k_B_T*, active force parameter 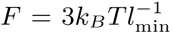, and the coupling parameter *β* = 20 *D/kBT*. (D) On the left, a schematic diagram of the vesicle where the curved proteins are coloured with red if it’s *x* coordinate *xi > x*CM, and with blue if *xi < x*CM. The centre of mass is denoted by a lime-coloured marker. On the right, the mean curvature for the curved proteins for *xi > x*CM and *xi < x*CM in red and blue respectively. The black dashed lines denote the times of the force reversals along the *x* direction.

In the SI section G.2 (Fig. S-9) we demonstrate these oscillations in a dumbbell pattern confined by adhesion (See Movie S-17), rather than rigid barriers, where the mechanism for the oscillations is identical: since the leading edge does not easily extend over the non-adhesive region, it loses its sharp edge, flattens and breaks up. We also demonstrate these oscillations in a simpler rectangular adhesive confinement (SI section H, See Movie S-18). In this case, the cell can get stuck for longer times at the end of the rectangle, as there is less space available for the leading edge to quickly break up into two opposing parts compared to the dumbbell pattern. Indeed, cells inside confining adhesion patterns often exhibit random oscillatory dynamics in experiments [45, 46], sometimes getting stuck at the ends of the patterns before switching their direction of migration [47].

Our model is based on a physical mechanism that inhibits the leading edge at the pattern’s edge, namely its curvature sensitivity, which explains both the behavior for adhesive and topographic confinement. Additional biochemical feedbacks may also contribute [9, 48]. Our results are also relevant to cellular oscillations observed when cells form their own adhesive “confinement” by de-position of extracellular matrix (ECM) [49, 50].

## IV. DISCUSSION AND CONCLUSION

Recently we have demonstrated that the coupling of curvature (through CMC) and recruitment of active protrusive forces due to actin polymerization, together with surface adhesion [12], provides a powerful organizing principle that can explain a variety of cellular shape dynamics and migration patterns [13, 14, 28]. Here we extended this model by implementing a simplified mechanism that couples the membrane organization of the CMC to an internal net actin flow that induces a polarity cue gradient across the cell. This polarity cue in turn introduces long-range inhibition of the local forces exerted by the CMC, thereby completing the feedback between the CMC organization and global polarization of the cell [25, 26].

This extension greatly increases the robustness of the polarized vesicle in our model. It allows us to use our model to explain a large variety of cellular dynamics which are observed in living cells, such as the relation between cell speed and aspect-ratio, cell-barrier scattering, and cellular oscillations in different types of geometric confinements. The agreement between the experiments and the model emphasizes that curved protein complexes are crucial in the formation and dynamics of lamellipodia-driven cell migration [18], and explain the sensitivity of the lamellipodium’s stability to its leading-edge curvature [51]. We demonstrate that simple, and therefore general (not cell-type specific), physics-based mechanisms play essential roles in directing cellular shape and migration. Biological and biochemical complexity allows cells to exert more precise control over these physical mechanisms, in response to different external conditions.

## V. ACKNOWLEDGMENTS

N.S.G. is the incumbent of the Lee and William Abramowitz Professorial Chair of Biophysics, and acknowledges support by the Israel Science Foundation (Grant No. 207/22). This research is made possible in part by the historic generosity of the Harold Perlman Family. The research of CB and C.M.-T. has been partially funded by the Deutsche Forschungsgemeinschaft (DFG), Project-ID No. 318763901–SFB1294

## Author contributions

SS and NSG designed the computational research; SS implemented the code and ran the simulations and data analysis; CMT and CB designed the experimental research; CMT performed the experiments; SP and AI developed the original code; SS and NSG wrote the paper, edited by CMT, CB, AI. All authors contributed to the article and approved the submitted version.

## Author declaration

The authors declare no competing interest.

---

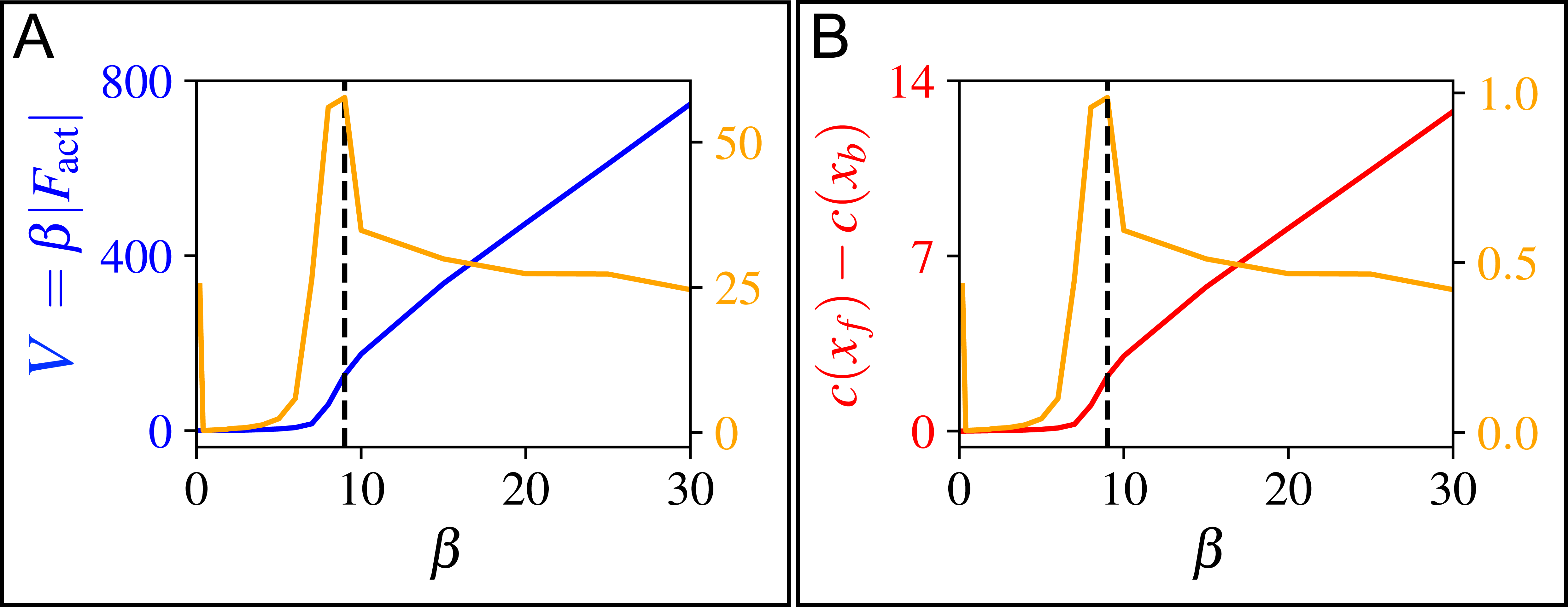

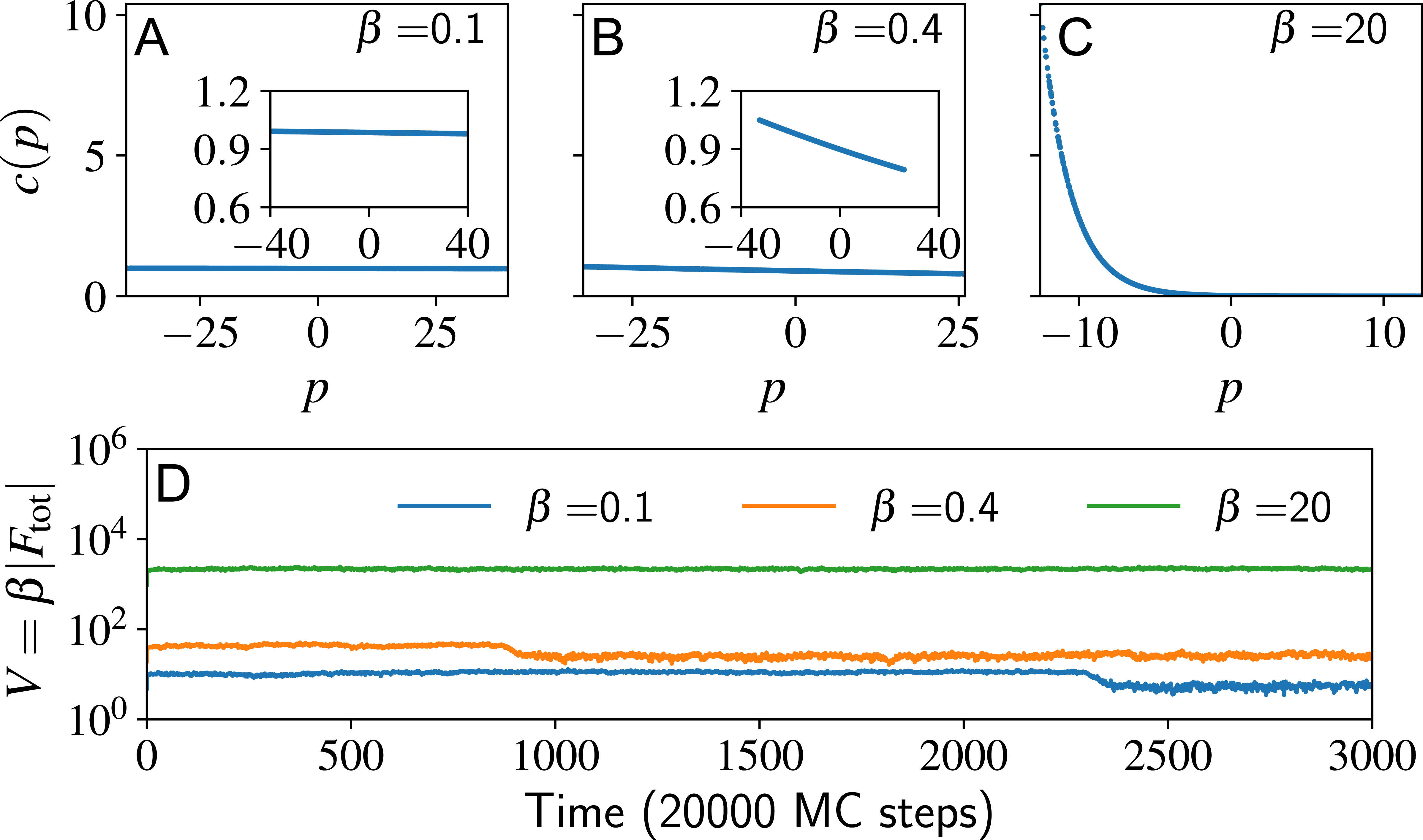

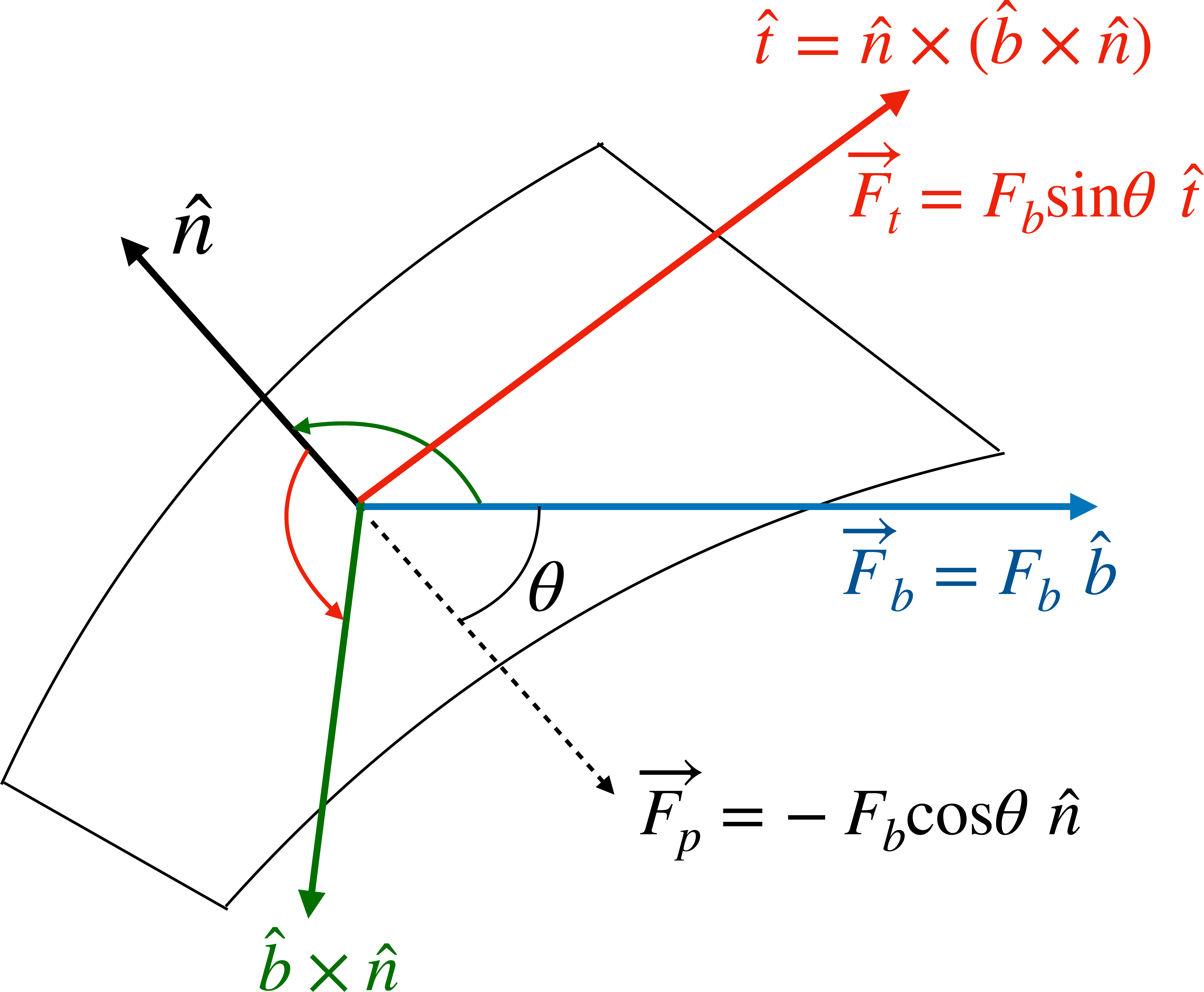

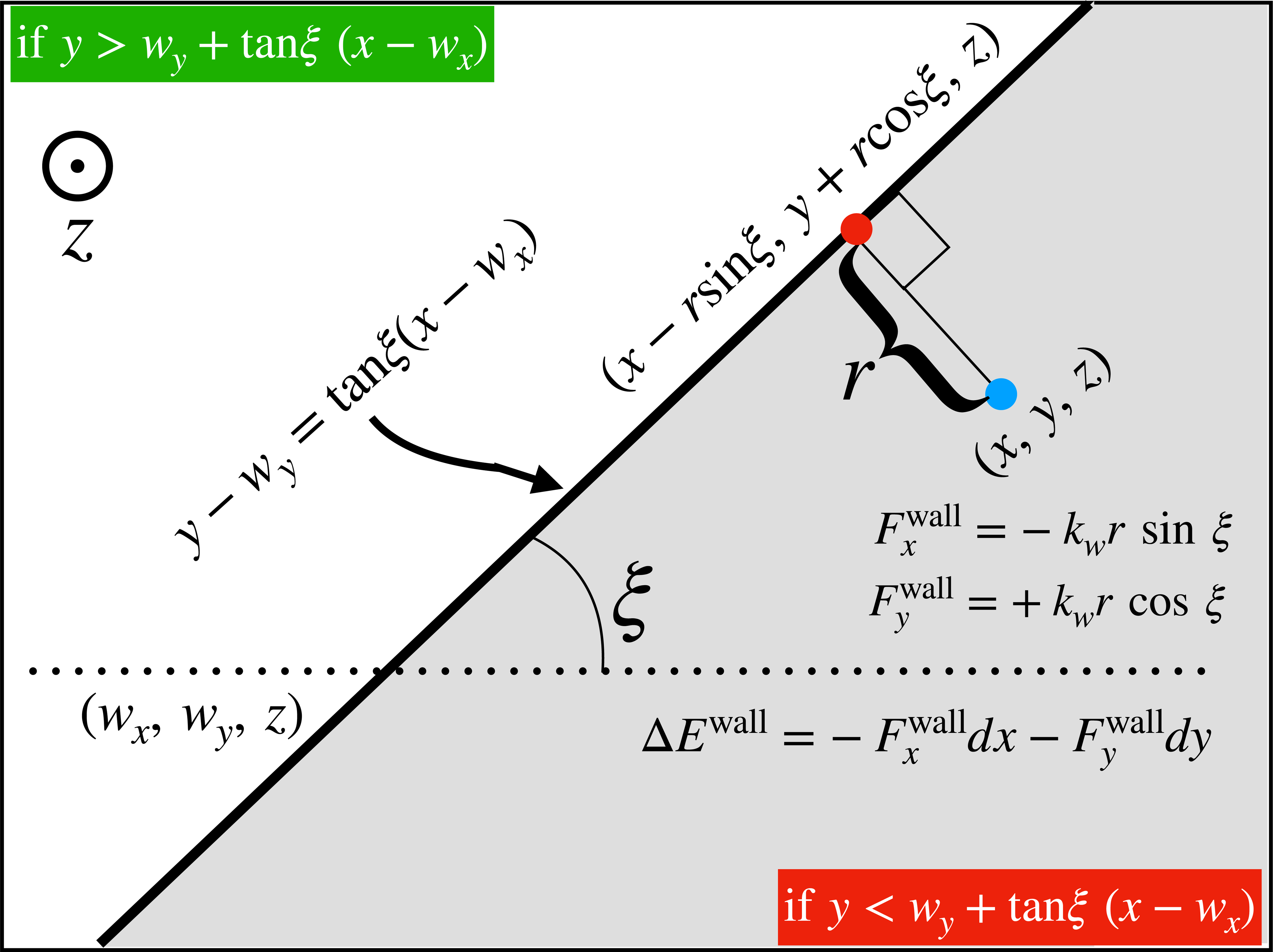

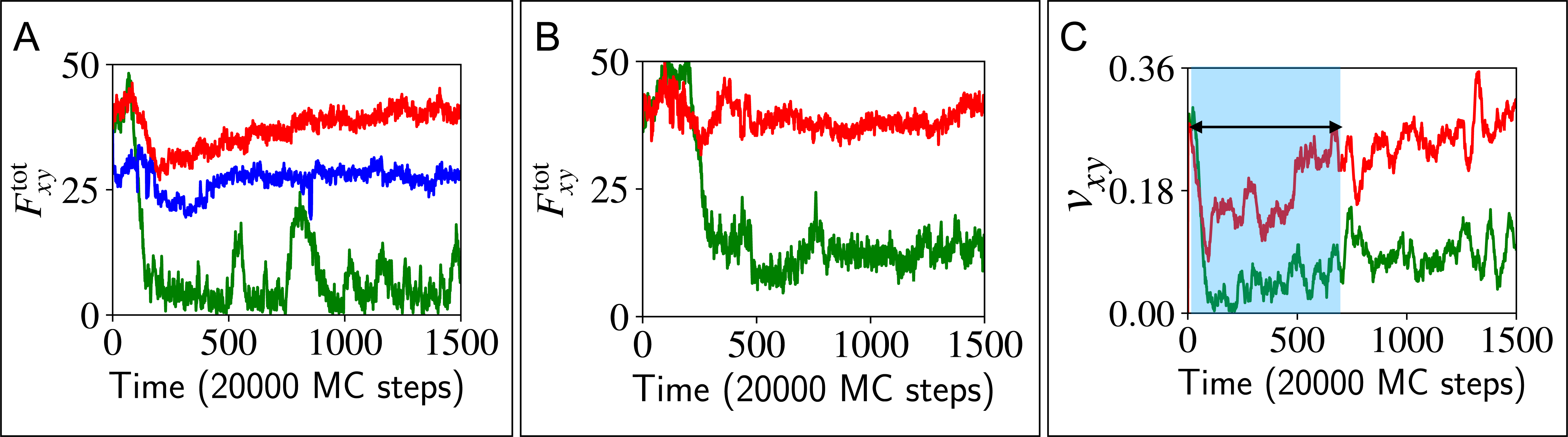

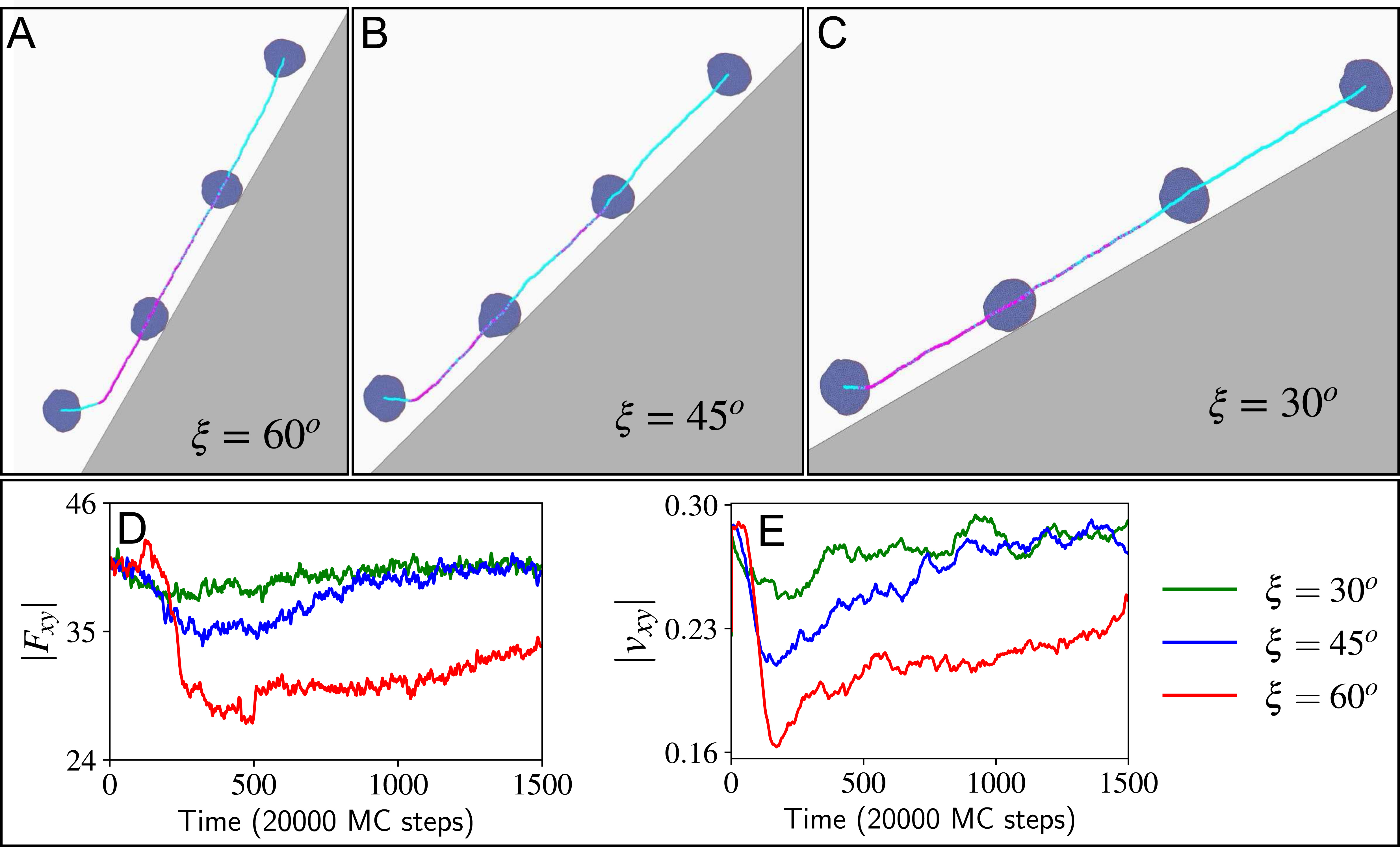

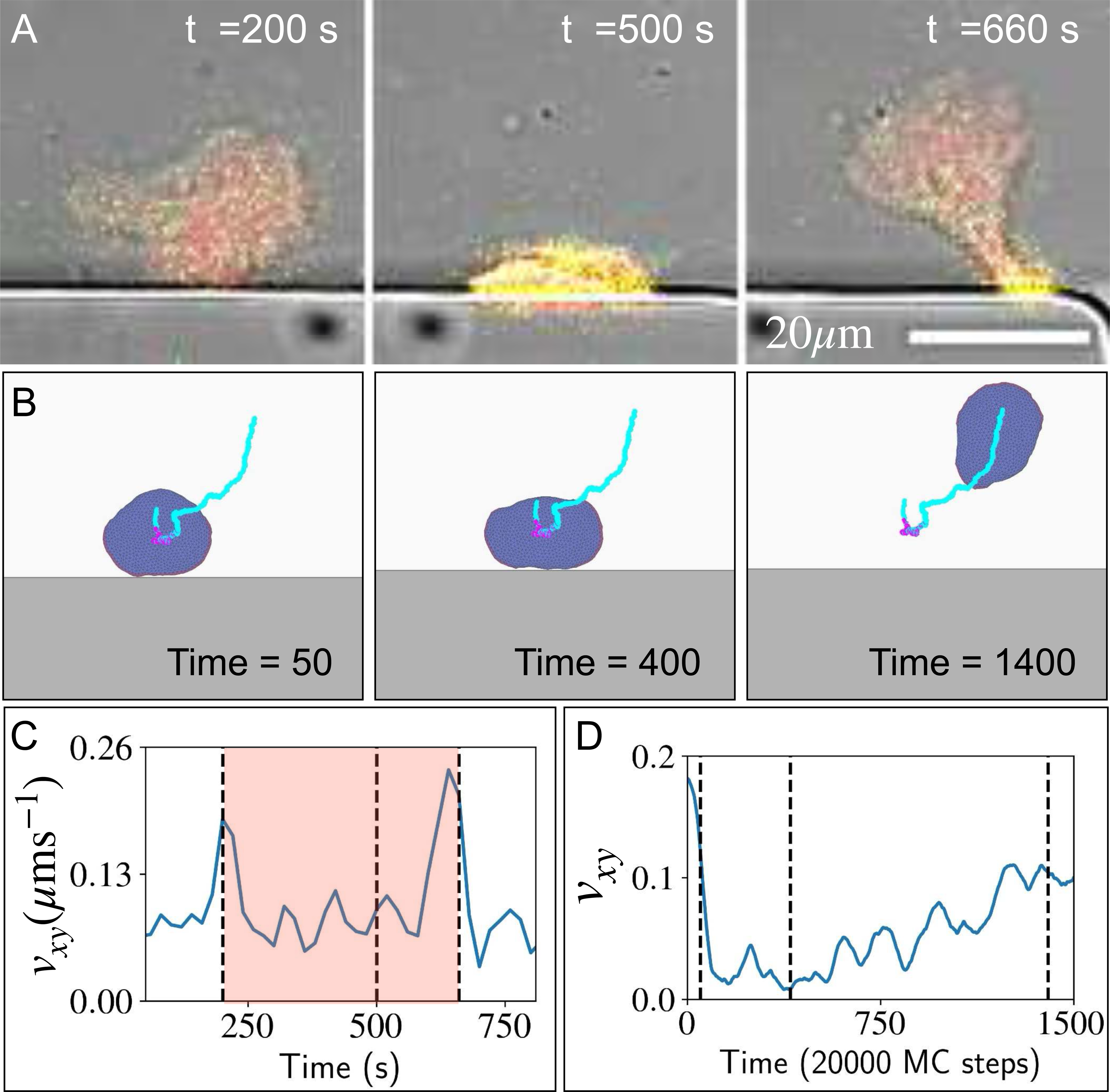

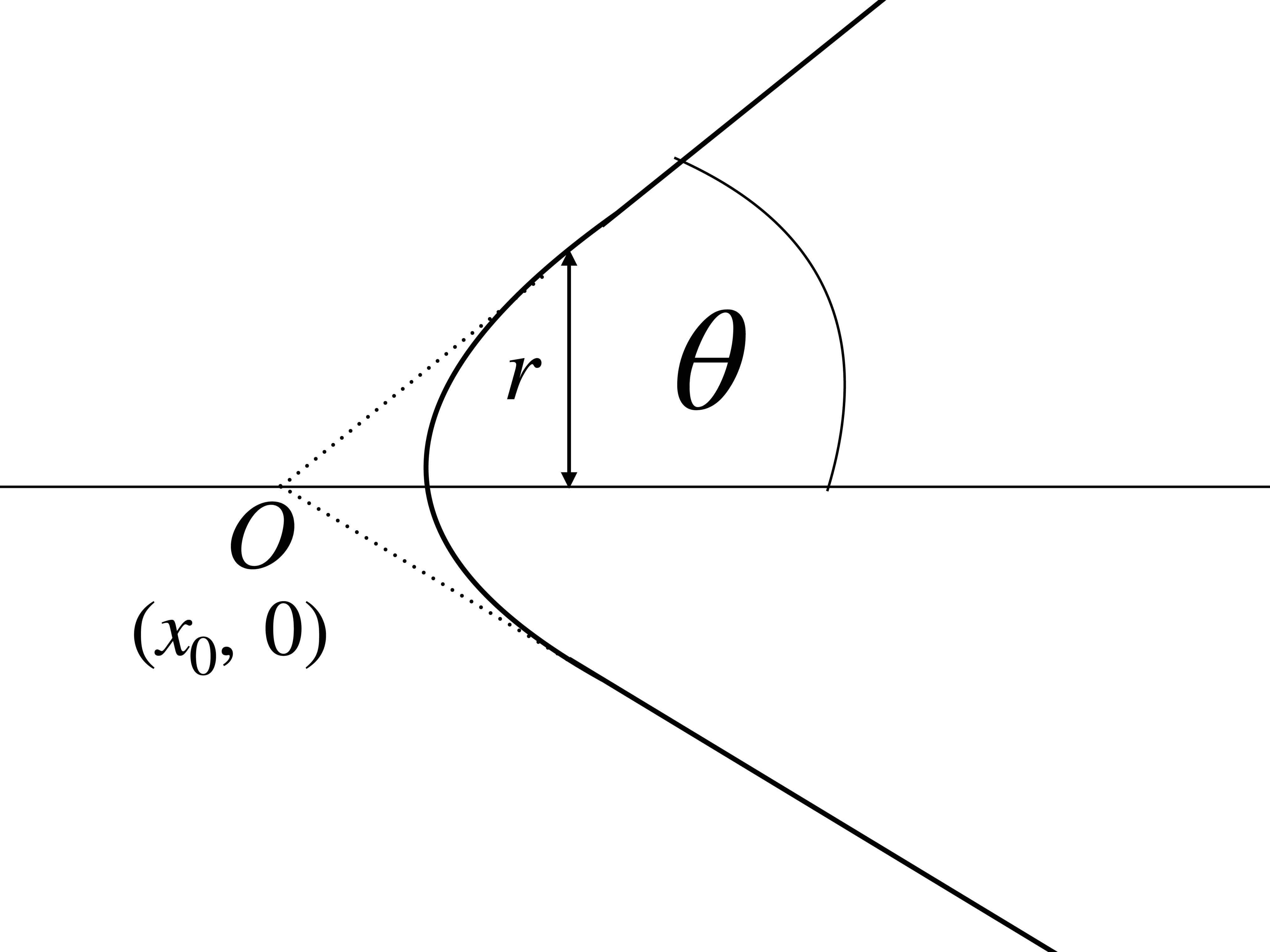

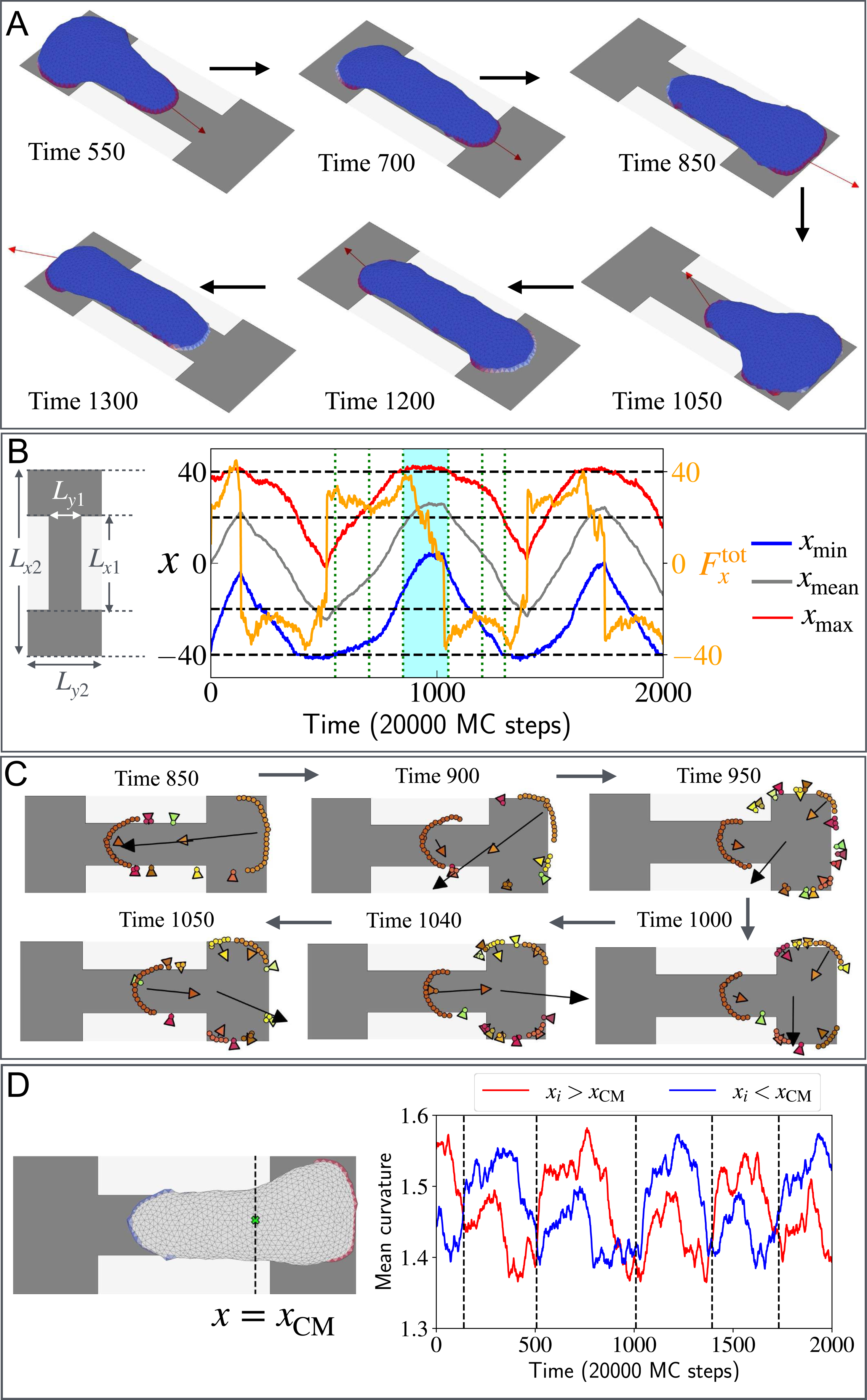

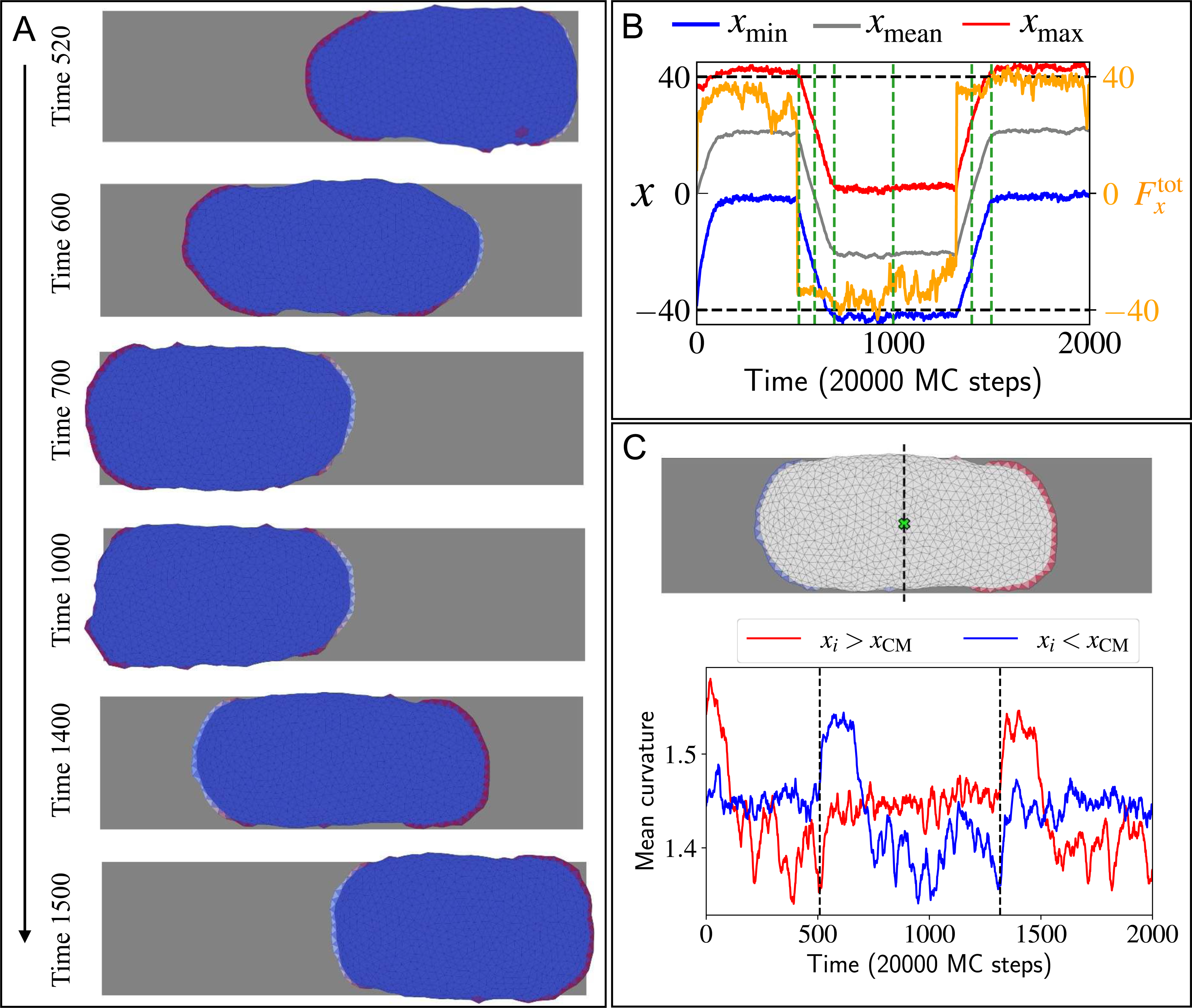

## References

[1] K. M. Yamada and M. Sixt, Mechanisms of 3d cell migration, Nature Reviews molecular cell biology 20, 738 (2019).

[2] Y.-G. Ko, C. C. Co, and C.-C. Ho, Directing cell migration in continuous microchannels by topographical amplification of natural directional persistence, Biomaterials 34, 353 (2013).

[3] M. K. Driscoll, X. Sun, C. Guven, J. T. Fourkas, and W. Losert, Cellular contact guidance through dynamic sensing of nanotopography, ACS nano 8, 3546 (2014).

[4] B. R. Graziano, J. P. Town, E. Sitarska, T. L. Nagy, M. Fošnarič, S. Penič, A. Iglič, V. Kralj-Iglič, N. S. Gov, A. Diz-Muñoz, et al., Cell confinement reveals a branched-actin independent circuit for neutrophil polarity, PLoS biology 17, e3000457 (2019).

[5] M. Werner, A. Petersen, N. A. Kurniawan, and C. V. Bouten, Cell-perceived substrate curvature dynamically coordinates the direction, speed, and persistence of stromal cell migration, Advanced Biosystems 3, 1900080 (2019).

[6] O. Nagel, C. Guven, M. Theves, M. Driscoll, W. Losert, and C. Beta, Geometry-driven polarity in motile amoeboid cells, PLoS ONE 9, e113382 (2014).

[7] M.-C. Kim, D. M. Neal, R. D. Kamm, and H. H. Asada, Dynamic modeling of cell migration and spreading behaviors on fibronectin coated planar substrates and micropatterned geometries, PLoS computational biology 9, e1002926 (2013).

[8] X. He and Y. Jiang, Substrate curvature regulates cell migration, Physical biology 14, 035006 (2017).

[9] B. A. Camley, Y. Zhao, B. Li, H. Levine, and W.-J. Rappel, Periodic migration in a physical model of cells on micropatterns, Physical review letters 111, 158102 (2013).

[10] B. Winkler, I. S. Aranson, and F. Ziebert, Confinement and substrate topography control cell migration in a 3d computational model, Communications Physics 2, 82 (2019).

[11] J. E. Ron, M. Crestani, J. M. Kux, J. Liu, N. Al-Dam, P. Monzo, N. C. Gauthier, P. J. Śaez, and N. S. Gov, Emergent seesaw oscillations during cellular directional decision-making, Nature Physics 20, 1 (2024).

[12] R. K. Sadhu, S. Penič, A. Iglič, and N. S. Gov, Modelling cellular spreading and emergence of motility in the presence of curved membrane proteins and active cytoskeleton forces, The European Physical Journal Plus 136, 495 (2021).

[13] R. K. Sadhu, A. Iglič, and N. S. Gov, A minimal cell model for lamellipodia-based cellular dynamics and migration, Journal of Cell Science 136, jcs260744 (2023).

[14] R. K. Sadhu, M. Luciano, W. Xi, C. Martinez-Torres, M. Schröder, C. Blum, M. Tarantola, S. Villa, S. Penič, A. Iglič, et al., A minimal physical model for curvotaxis driven by curved protein complexes at the cell’s leading edge, Proceedings of the National Academy of Sciences 121, e2306818121 (2024).

[15] J. Linkner, G. Witte, H. Zhao, A. Junemann, B. Nordholz, P. Runge-Wollmann, P. Lappalainen, and J. Faix, The inverse bar domain protein ibara drives membrane remodeling to control osmoregulation, phagocytosis and cytokinesis, Journal of Cell Science 127, 1279 (2014).

[16] I. Begemann, T. Saha, L. Lamparter, I. Rathmann, D. Grill, L. Golbach, C. Rasch, U. Keller, B. Trappmann, M. Matis, et al., Mechanochemical self-organization determines search pattern in migratory cells, Nature Physics 15, 848 (2019).

[17] A. Pipathsouk, R. M. Brunetti, J. P. Town, B. R. Graziano, A. Breuer, P. A. Pellett, K. Marchuk, N.-H. T. Tran, M. F. Krummel, D. Stamou, et al., The wave complex associates with sites of saddle membrane curvature, Journal of Cell Biology 220, e202003086 (2021).

[18] M. Wu, P. Marchando, K. Meyer, Z. Tang, D. N. Woolfson, and O. D. Weiner, The wave complex forms linear arrays at negative membrane curvature to instruct lamellipodia formation., bioRxiv, 2024 (2024).

[19] L. Baldauf, F. Frey, M. Arribas Perez, T. Idema, and G. H. Koenderink, Branched actin cortices reconstituted in vesicles sense membrane curvature, Biophysical Journal 122, 2311–2324 (2023).

[20] N. Andrew and R. H. Insall, Chemotaxis in shallow gradients is mediated independently of ptdins 3-kinase by biased choices between random protrusions, Nature cell biology 9, 193 (2007).

[21] T. D. Yang, J.-S. Park, Y. Choi, W. Choi, T.-W. Ko, and K. J. Lee, Zigzag turning preference of freely crawling cells, PLoS One 6, e20255 (2011).

[22] P. Dieterich, R. Klages, R. Preuss, and A. Schwab, Anomalous dynamics of cell migration, Proceedings of the National Academy of Sciences 105, 459 (2008).

[23] O. D. Weiner, W. A. Marganski, L. F. Wu, S. J. Altschuler, and M. W. Kirschner, An actin-based wave generator organizes cell motility, PLoS biology 5, e221 (2007).

[24] S. Gross-Thebing, L. Truszkowski, D. Tenbrinck, H. Śanchez-Iranzo, C. Camelo, K. J. Westerich, A. Singh, P. Maier, J. Prengel, P. Lange, et al., Using migrating cells as probes to illuminate features in live embryonic tissues, Science Advances 6, eabc5546 (2020).

[25] P. Maiuri, J.-F. Rupprecht, S. Wieser, V. Ruprecht, O. Bénichou, N. Carpi, M. Coppey, S. De Beco, N. Gov, C.-P. Heisenberg, et al., Actin flows mediate a universal coupling between cell speed and cell persistence, Cell 161, 374 (2015).

[26] J. E. Ron, P. Monzo, N. C. Gauthier, R. Voituriez, and N. S. Gov, One-dimensional cell motility patterns, Physical review research 2, 033237 (2020).

[27] M. Fošnarič, S. Penič, A. Iglič, V. Kralj-Iglič, M. Drab, and N. S. Gov, Theoretical study of vesicle shapes driven by coupling curved proteins and active cytoskeletal forces, Soft Matter 15, 5319 (2019).

[28] S. Sadhukhan, S. Penič, A. Iglič, and N. S. Gov, Modelling how curved active proteins and shear flow pattern cellular shape and motility, Frontiers in Cell and Developmental Biology 11, 1193793 (2023).

[29] I. Lavi, N. Meunier, R. Voituriez, and J. Casademunt, Motility and morphodynamics of confined cells, Physical Review E 101, 022404 (2020).

[30] O. D. Weiner, Regulation of cell polarity during eukaryotic chemotaxis: the chemotactic compass, Current opinion in cell biology 14, 196 (2002).

[31] Y. Mori, A. Jilkine, and L. Edelstein-Keshet, Wave-pinning and cell polarity from a bistable reaction-diffusion system, Biophysical journal 94, 3684 (2008).

[32] A. R. Houk, A. Jilkine, C. O. Mejean, R. Boltyanskiy, E. R. Dufresne, S. B. Angenent, S. J. Altschuler, L. F. Wu, and O. D. Weiner, Membrane tension maintains cell polarity by confining signals to the leading edge during neutrophil migration, Cell 148, 175 (2012).

[33] N. W. Goehring and S. W. Grill, Cell polarity: mechanochemical patterning, Trends in cell biology 23, 72 (2013).

[34] W.-J. Rappel and L. Edelstein-Keshet, Mechanisms of cell polarization, Current Opinion in Systems Biology 3, 43 (2017), • Mathematical modelling • Mathematical modelling, Dynamics of brain activity at the systems level • Clinical and translational systems biology.

[35] R. Illukkumbura, T. Bland, and N. W. Goehring, Patterning and polarization of cells by intracellular flows, Current Opinion in Cell Biology 62, 123 (2020), cell Architecture.

[36] T. Moldenhawer, E. Moreno, D. Schindler, S. Flemming, M. Holschneider, W. Huisinga, S. Alonso, and C. Beta, Spontaneous transitions between amoeboid and keratocyte-like modes of migration, Frontiers in Cell and Developmental Biology 10, 10.3389/fcell.2022.898351 (2022).

[37] A. B. Verkhovsky, T. M. Svitkina, and G. G. Borisy, Self-polarization and directional motility of cytoplasm, Current Biology 9, 11 (1999).

[38] K. Keren, Z. Pincus, G. M. Allen, E. L. Barnhart, G. Marriott, A. Mogilner, and J. A. Theriot, Mechanism of shape determination in motile cells, Nature 453, 475 (2008).

[39] F. Raynaud, M. E. Ambühl, C. Gabella, A. Bornert, I. F. Sbalzarini, J.-J. Meister, and A. B. Verkhovsky, Minimal model for spontaneous cell polarization and edge activity in oscillating, rotating and migrating cells, Nature Physics 12, 367 (2016).

[40] E. Ghabache, Y. Cao, Y. Miao, A. Groisman, P. N. Devreotes, and W. Rappel, Coupling traction force patterns and actomyosin wave dynamics reveals mechanics of cell motion, Molecular Systems Biology 17, 10.15252/msb.202110505 (2021).

[41] E. Sitarska, S. D. Almeida, M. S. Beckwith, J. Stopp, J. Czuchnowski, M. Siggel, R. Roessner, A. Tschanz, C. Ejsing, Y. Schwab, et al., Sensing their plasma membrane curvature allows migrating cells to circumvent obstacles, Nature Communications 14, 5644 (2023).

[42] A. Roycroft and R. Mayor, Molecular basis of contact inhibition of locomotion, Cellular and Molecular Life Sciences 73, 1119 (2016).

[43] L. Truszkowski, D. Batur, H. Long, K. Tarbashevich, B. E. Vos, B. Trappmann, and E. Raz, Primordial germ cells adjust their protrusion type while migrating in different tissue contexts in vivo, Development 150, dev200603 (2023).

[44] F. A. Gegenfurtner, B. Jahn, H. Wagner, C. Ziegenhain, W. Enard, L. Geistlinger, J. O. Rädler, A. M. Vollmar, and S. Zahler, Micropatterning as a tool to identify regulatory triggers and kinetics of actin-mediated endothelial mechanosensing, Journal of Cell Science 131, jcs212886 (2018).

[45] A. Fink, D. B. Brückner, C. Schreiber, P. J. Röttgermann, C. P. Broedersz, and J. O. Rädler, Area and geometry dependence of cell migration in asymmetric two-state micropatterns, Biophysical journal 118, 552 (2020).

[46] Y. Kalukula, M. Luciano, G. Charras, D. Brueckner, and S. Gabriele, The actin cortex acts as a mechanical memory of morphology in confined migrating cells, bioRxiv, 2024 (2024).

[47] F. Zhou, S. A. Schaffer, C. Schreiber, F. J. Segerer, A. Goychuk, E. Frey, and J. O. Radler, Quasi-periodic migration of single cells on short microlanes, PLoS One 15, e0230679 (2020).

[48] P. Zadeh and B. A. Camley, Nonlinear dynamics of confined cell migration – modeling and inference, arXiv 10.48550/ARXIV.2404.07390 (2024).

[49] J. d’Alessandro, A. Barbier-Chebbah, V. Cellerin, O. Benichou, R. M. Mège, R. Voituriez, and B. Ladoux, Cell migration guided by long-lived spatial memory, Nature Communications 12, 4118 (2021).

[50] E. Perez Ipiña, J. d’Alessandro, B. Ladoux, and B. A. Camley, Deposited footprints let cells switch between confined, oscillatory, and exploratory migration, Proceedings of the National Academy of Sciences 121, e2318248121 (2024).

[51] C. Jiang, H.-Y. Luo, X. Xu, S.-X. Dou, W. Li, D. Guan, F. Ye, X. Chen, M. Guo, P.-Y. Wang, et al., Switch of cell migration modes orchestrated by changes of three-dimensional lamellipodium structure and intracellular diffusion, Nature Communications 14, 5166 (2023).

