## Supplementary material for "Modelling how lamellipodia-driven cells maintain persistent migration and interact with external barriers"

Shubhadeep Sadhukhan* and Nir Gov†
*Department of Chemical and Biological Physics, Weizmann Institute of Science, Rehovot, Israel*

Cristina Martinez-Torres‡
*Institute of Physics and Astronomy, University of Potsdam, Potsdam 14476, Germany*

Samo Penič§
*Laboratory of Physics, Faculty of Electrical Engineering, University of Ljubljana, Ljubljana, Slovenia*

Carsten Beta¶
*Institute of Physics and Astronomy, University of Potsdam, Potsdam 14476, Germany and Nano Life Science Institute (WPI-NanoLSI), Kanazawa University, Kanazawa 920-1192, Japan*

Aleš Iglič**
*Laboratory of Physics, Faculty of Electrical Engineering, University of Ljubljana, Ljubljana, Slovenia and Laboratory of Clinical Biophysics, Faculty of Medicine, University of Ljubljana, Ljubljana, Slovenia*

(Dated: September 6, 2024)

### MATERIALS AND METHODS

#### A. Theoretical model

We modelled the cell membrane as a three-dimensional vesicle which is described by a closed surface with $N$ vertices connected to its neighbours by bonds and it forms a dynamically triangulated, self-avoiding network, with the topology of sphere [5, 6]. $\boldsymbol{r_i}$ is the position vector of the $i$th vertex. All the lengths are measured in a scale of $l_{\text{min}}$. There is a percentage $\rho = 100N_c/N$ of vertex sites that represent the proteins that induce cytoskeletal active forces. The vesicle is placed on a uniform adhesive substrate. The vesicle energy has four components: The bending energy is given by,

$$W_b = \frac{\kappa}{2} \int_A (C_1 + C_2 - C_0)^2 \; dA, \tag{S-1}$$

where, $\kappa$ is the bending rigidity, $C_1$, $C_2$ are the principal curvatures and $C_0$ is the spontaneous curvature. We consider the spontaneous curvature $C_0 = 1l_{\text{min}}^{-1}$ for the curved protein sites represented in red and blue represents the bare membrane for which $C_0 = 0$. We set $\kappa = 20k_BT$ throughout the paper except for the cases we mentioned. The protein-protein interaction energy is given by,

$$W_d = -w \sum_{i<j} \mathcal{H}(r_0 - r_{ij}) \tag{S-2}$$

where $\mathcal{H}$ is the Heaviside function, $r_{ij} = |\boldsymbol{r_i} - \boldsymbol{r_j}|$ is the distance between protein sites, $r_0$ is the range interaction and $w$ is the strength. We set the parameter $w$ to $1k_BT$ throughout the paper. The energy due to the active force is given by,

$$W_F = -F \sum_i \hat{n}_i \cdot \boldsymbol{r_i} \tag{S-3}$$

*
†
‡
§
¶
**

where $F$ is the magnitude of the active force, $\hat{n}_i$ is the outward unit normal vector of the $i$th protein site vertex and $\boldsymbol{r_i}$ is the position vector of the protein. Finally, the adhesion energy due to the extra-cellular substrate is given by,

$$W_A = -\sum_{i'} E_{\mathrm{ad}} \tag{S-4}$$

where $E_{\mathrm{ad}}$ is the adhesion strength, and the sum runs over all the adhered vertices to the substrate i.e., the $z$ component of the vertex position is within a range $z_{\mathrm{ad}} < z_i < z_{\mathrm{ad}} + \delta z$. We set $\delta z = 1 l_{\mathrm{min}}$ throughout the paper.

### B. Calculation of direction of net actin flow

For simplicity, we want to map the whole three-dimensional problem into a simplified one-dimensional model as the concentration profile of the inhibiting molecules is calculated for one-dimensional case [4]. We want to approximate the direction along which we want to calculate the variation of the concentration of inhibitors. First, curved-membrane protein (CMP) components on the vesicle membrane are clusterized. Let us consider the $i$th cluster that consists of $N_i$ number of proteins. The positions of the $j$th protein-vertex of the $i$th cluster is $(x^i_j,\ y^i_j,\ z^i_j)$. We approximated the actin flow from the protein site towards the centre of mass of the vesicle $\boldsymbol{r}_{\mathrm{CM}}(x_{\mathrm{CM}},\ y_{\mathrm{CM}},\ z_{\mathrm{CM}})$ and the magnitude of the flow is proportional to the active force. We compute the centre of mass of the vesicle by taking the average position of all vertices on the membrane. Therefore, we find a vector $\boldsymbol{c}^i{=}(c^i_x,\ c^i_y,\ c^i_z)$ for the $i$th cluster by,

$$\begin{aligned} c^i_x &= \frac{1}{N_i}\sum_j F^i_j (x^i_j - x_{\mathrm{CM}}) \\ c^i_y &= \frac{1}{N_i}\sum_j F^i_j (y^i_j - y_{\mathrm{CM}}) \\ c^i_z &= \frac{1}{N_i}\sum_j F^i_j (z^i_j - z_{\mathrm{CM}}) \end{aligned} \tag{S-5}$$

where, $j$ runs over all the vertices within the $i$th cluster. $F^i_j$ is the magnitude of the active force by the $j$th vertex in the $i$th cluster. For the direction of net actin flow, we fit a three-dimensional line through the centre of mass and the $\boldsymbol{c}^i$ vectors for all the clusters with a weightage proportional to their size $N_i$.

Next, we find the symmetric covariance matrix as follows,

$$C = \begin{bmatrix} a_{xx} & a_{xy} & a_{xz} \\ a_{yx} & a_{yy} & a_{yz} \\ a_{zx} & a_{zy} & a_{zz} \end{bmatrix}. \tag{S-6}$$

where the elements of this matrix are given by,

$$\begin{aligned} a_{xx} &= \sum_i N_i (c^i_x - x_{\mathrm{CM}})^2 \\ a_{yy} &= \sum_i N_i (c^i_y - y_{\mathrm{CM}})^2 \\ a_{zz} &= \sum_i N_i (c^i_z - z_{\mathrm{CM}})^2 \\ a_{xy} = a_{yx} &= \sum_i N_i (c^i_x - x_{\mathrm{CM}})(c^i_y - y_{\mathrm{CM}}) \\ a_{yz} = a_{zy} &= \sum_i N_i (c^i_y - y_{\mathrm{CM}})(c^i_z - z_{\mathrm{CM}}) \\ a_{zx} = a_{xz} &= \sum_i N_i (c^i_z - z_{\mathrm{CM}})(c^i_x - x_{\mathrm{CM}}). \end{aligned} \tag{S-7}$$

Then, we find the eigenvector $\hat{\boldsymbol{e}}$ corresponding to the highest eigenvalue using the "power method" with a tolerance of $10^{-3}$. This method gives the direction of net actin flow with an inaccuracy up to a sign reversal. To correct this we use a physical condition that the total active force and the net actin flow are opposite to each other, i.e., $\boldsymbol{F}^{\mathbf{tot}} \cdot \hat{\boldsymbol{e}} < 0$.

#### C. UCSP time step

First, we find the axis of the net actin flow using the cluster information. Now, we find the projection of each vertex position with respect to the centre of mass of the vesicle on the computed flow axis. This projection for the $i$th vertex is given by,

$$p_i = -(\boldsymbol{r}_i - \boldsymbol{r}_{\mathrm{CM}}) \cdot \hat{\boldsymbol{e}}. \tag{S-8}$$

The negative sign is for the convention so the maximum projection $p_{\max}$ and $p_{\min}$ denote the front and rear of the vesicle. From this projection, we compute the concentration of the polarity cue component, and hence we get the protrusive active force $\tilde{F}$ (Eq. 3equation.2.3) including the inhibition effect. As the total amount of the polarity cue is constant $c_{\mathrm{tot}}$, we need to modify the factor in the concentration profile of the polarity cue given in previous work [4] for one-dimension. This factor is modified due to the fact we have a three-dimensional vesicle and therefore we need to integrate over the whole vesicle volume even though the variation of the approximated net actin flow is in one direction. Therefore, an additional factor $(p_{\max} - p_{\min})/V_{\mathrm{ves}}$ get multiplied to the one-dimensional form given in [4]. Here, $V_{\mathrm{ves}}$ is the volume of the vesicle and $p_{\max} - p_{\min}$ denotes the elongation of the vesicle in the direction of the net actin flow. The complete updated form is given in the Eq. 2equation.2.2. We repeat this polarity cue concentration calculation until the net actin flow direction converges to a given tolerance level of 0.1%. If the change of direction of net actin flow due to the shape change of the vesicle is more than 10% we repeat the calculation of the concentration of polarity cues. If $\boldsymbol{e}$=($e_1$, $e_2$, $e_3$) is the net actin flow at the step when we computed the concentration profile and $\boldsymbol{e'}$=($e'_1$, $e'_2$, $e'_3$) is the of net actin flow after the change in vesicle shape, we find the relative change in a vector by,

$$\text{error} = \frac{(e'_1 - e_1)^2 + (e'_2 - e_2)^2 + (e'_3 - e_3)^2}{e_1^2 + e_2^2 + e_3^2}. \tag{S-9}$$

If the error becomes greater than 0.1, i.e., an error of 10% we repeat another UCSP calculation.

| **Important Parameters List** | | |
|---|---|---|
| Number of vertices | $N$ | 1447 |
| Bending rigidity | $\kappa$ | $20k_BT$ |
| Protein-protein interaction energy | $w$ | $1k_BT$ |
| Intrinsic curvature of proteins | $C_0$ | $1\ l_{\mathrm{min}}^{-1}$ |
| Range of adhesion | $\delta z$ | $1l_{\mathrm{min}}$ |
| $z$ coordinate of adhesion plane | $z_{\mathrm{ad}}$ | $-10.725l_{\mathrm{min}}$ |
| Diffusion coefficient of inhibitors | $D$ | 4000 |
| Total amount of inhibitors | $c_{\mathrm{tot}}$ | 4000 |
| Wall position $x$ coordinate | $w_x$ | $25l_{\mathrm{min}}$ |
| Wall position $y$ coordinate | $w_y$ | $0l_{\mathrm{min}}$ |
| Spring constant of the wall | $k_w$ | $0.1k_BTl_{\mathrm{min}}^{-2}$ |
| Smoothness parameter for the triangular tip | $r$ | $5l_{\mathrm{min}}$ |
| Asymptotic angle for the triangular tip | $\theta$ | $45^o$ |
| Peak force for blowing from behind | $F_{\mathrm{blow}}$ | $20k_BTl_{\mathrm{min}}^{-1}$ |
| Range for blowing from behind | $\Delta x$ | $2l_{\mathrm{min}}$ |
| Rectangular shaped adhesive pattern | $L_x$<br>$L_y$ | 80 $l_{\mathrm{min}}$<br>22 $l_{\mathrm{min}}$ |
| Dumbbell shaped adhesive pattern | $L_{x1}$<br>$L_{x2}$<br>$L_{y1}$<br>$L_{y2}$ | 40 $l_{\mathrm{min}}$<br>80 $l_{\mathrm{min}}$<br>14 $l_{\mathrm{min}}$<br>32 $l_{\mathrm{min}}$ |
| Dumbbell shaped confinement box | $L_{x1}$<br>$L_{x2}$<br>$L_{y1}$<br>$L_{y2}$ | 36 $l_{\mathrm{min}}$<br>78 $l_{\mathrm{min}}$<br>16 $l_{\mathrm{min}}$<br>32 $l_{\mathrm{min}}$ |

TABLE I. All the important parameters used in the simulations

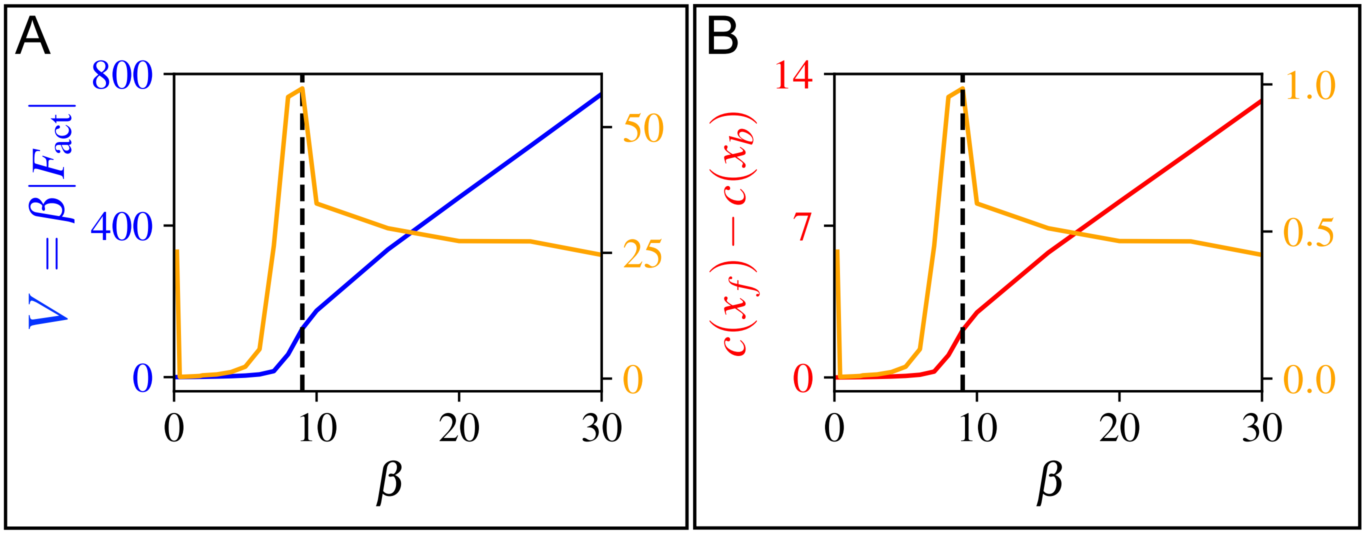


FIG. S-1. Demonstration of critical $\beta$: We turned off all the vertex movement and bond flips for this test. The orange lines in both panels show the first derivatives of the quantities net actin flow $V = \beta|F^{\rm tot}|$ and asymmetry in polarity cue concentration $c(p_f) - c(p_b)$ shown in the panel A and B respectively. The derivative shows a peak at the point of the critical coupling parameter denoted by a black dashed line. We used a two-arc-shaped vesicle which was spread on the adhesion $E_{\rm ad} = 1\ k_BT$ and $F = 3k_BT\ l_{\rm min}^{-1}$.

#### D. Breaking of protein cluster

We demonstrated that as the coupling between the asymmetry in polarity cues and the actin flow ($\beta$) increases, the UCSP mechanism can stabilize the crescent-shaped vesicle and hinder the transition to the non-motile shape (Fig. 2).We showed the concentration profile of the polarity cues over the "mini-cell" (vesicle) along the net actin flow axis for various coupling strength $\beta = 0.1,\ 0.4$, and 20 in units of $D/k_BT$ as shown in Fig. S-2 (A-C). When, $\beta$ is too small the concentration of such polarity cues is nearly constant. However, for the high value of $\beta$ can create a exponential fall of the concentration of polarity cues at the front of the "mini-cell". Next, we can see that time-series plot of the net actin flow $V = \beta|F^{\rm tot}|$ and it increases as coupling parameter $\beta$ increases as shown in Fig. S-2D.

#### E. Model of blowing force from behind

We model the force due to flowing fluid from the back of the vesicle. The force has the peak magnitude of $F_{\rm blow}$. It is acting on the vesicle from behind in the direction of $\hat{x}$. The blowing force acts only if the $x$ component of the outward normal is negative. This force magnitude also decays as a Gaussian-like function from its peak value $F_{\rm blow}$ at $(y_{\rm CM},\ z_{\rm ad})$ where $z_{\rm ad}$ is the $z$ coordinate of the adhesive substrate. Therefore, mathematically we can write this force magnitude at some point $(x, y, z)$ as,

$$F_b = F_{\rm blow} {\rm exp}\left( -\frac{1}{\sigma}\left[(y - y_{\rm CM})^2 + (z - z_{\rm ad})^2\right]\right), \tag{S-10}$$

where $\sigma = 100 l_{\rm min}^2$. Note that the blowing force is not varying with $x$ but it acts only from the back within a range from $x_{\rm back}$ to $x_{\rm back} + \Delta x$, where, $x_{\rm back}$ is the $x$ coordinate of the vesicle at the back and $\Delta x = 2l_{\rm min}$ is the range of the force. Now, this blowing force $F_b$ is decomposed into two parts, (a) Pressure part: $\boldsymbol{F_p} = -F_b \cos\theta\ \hat{n}$, (b) Tangential drag force: $\boldsymbol{F_t} = F_b \sin\theta\ \hat{t}$ where, $\hat{n}$ is the unit outward normal and $\hat{t}$ is the direction of tangential drag due to the blowing force from behind as shown in Fig. S-3.

#### F. Wall implementation

In our model, the vesicle is adhered to a plane $z = z_{\rm ad}$. Therefore, it moves on the $x - y$ plane. We implement the wall by drawing a line on $x - y$ plane and the wall is semi-infinitely extended in $z$ direction for the region $z \geq z_{\rm ad}$.

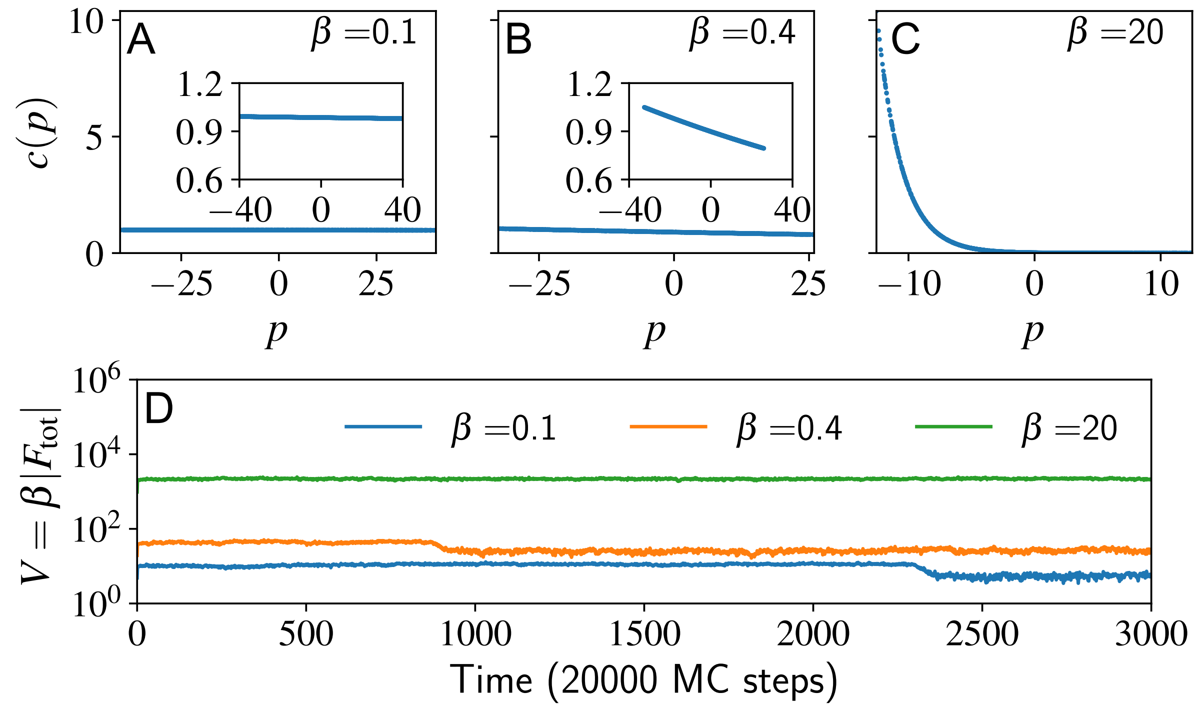


FIG. S-2. We showed the concentration profile along the UCSP axis at the final time 2999 in the units of 20000 MC steps for the cases $\beta = 0.1,\ 0.4$, and 20 in units of $D/k_BT$ respectively in (A-C). We can look into the concentration profile in a zoomed inset for the cases of $\beta = 0.1,\ 0.4\ D/k_BT$. We showed the time series of the magnitude of internal actin flow $V$ in D for different non-zero $\beta$.

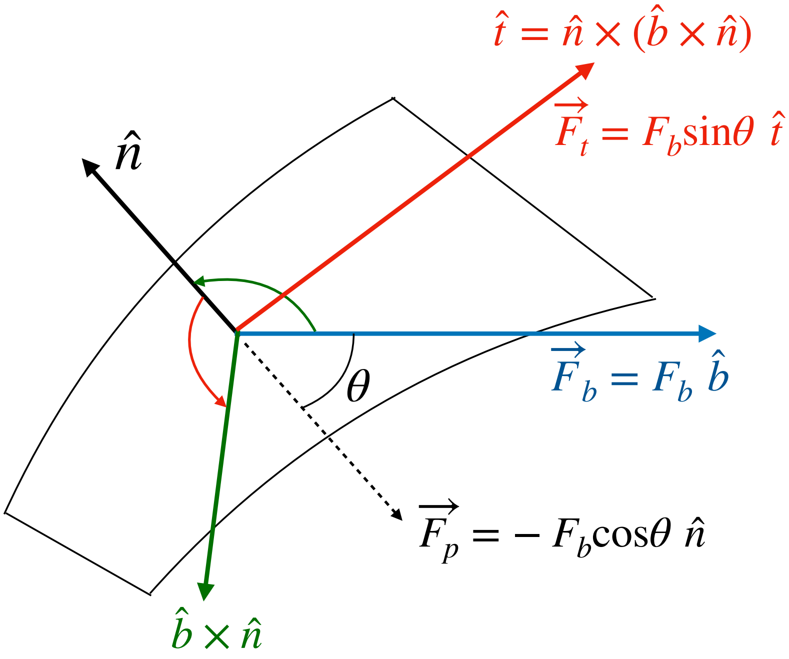


FIG. S-3. This force has two components $\boldsymbol{F_p} = -F_b \cos\theta\ \hat{n}$ and $\boldsymbol{F_t} = F_b \sin\theta\ \hat{t}$. The first one is the force due to pressure in the opposite direction of the outward normal of the membrane. The next one is the tangential drag which is tangential to the membrane. The figure shows the calculation of the directions.

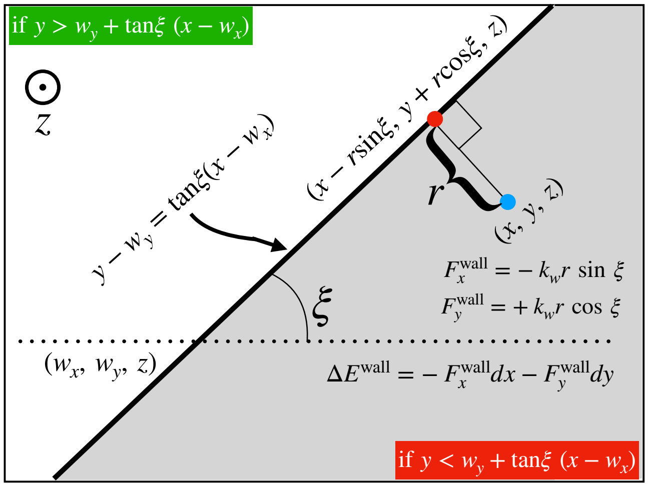


FIG. S-4. Schematic diagram of the wall implementation: The wall is specified by a straight line on $x - y$ plane which passes through the point $(w_x,\ w_y)$ and makes an angle $\xi$ with the $x$ axis. The wall is extended semi-infinitely in $z$ direction (Outward normal of the page) for the region $z \geq z_{\rm ad}$. We showed the region beyond the wall using a grey shade. Mathematically, if the condition $y < w_x + \tan\,\xi(x - w_x)$ holds, the vertex is trying to penetrate the wall. For a completely rigid wall ($k_w \to \infty$), the energy cost is infinite. Hence, no penetration is allowed. In the case of a soft wall, we model the softness of the wall by Hook's law where the vertex of the vesicle feels a restoring force depending on the position where the vertex is trying to penetrate. If $(x,\ y,\ z)$ is the position of the vertex, then the restoring force applied on it is given by, $(-k_w r\ \sin\xi,\ k_w r\ \cos\xi,\ 0)$, where $r = (x - w_x)\sin\xi - (y - w_x)\cos\xi$ is the perpendicular distance of the wall from the vertex when the vertex is in the grey shaded region. Therefore, if the vertex displaced $\boldsymbol{dr} = (dx,\ dy,\ dz)$ it costs an amount of energy due to the soft wall is $\Delta E^{\rm wall} = -F_x^{\rm wall}dx - F_y^{\rm wall}dy$.

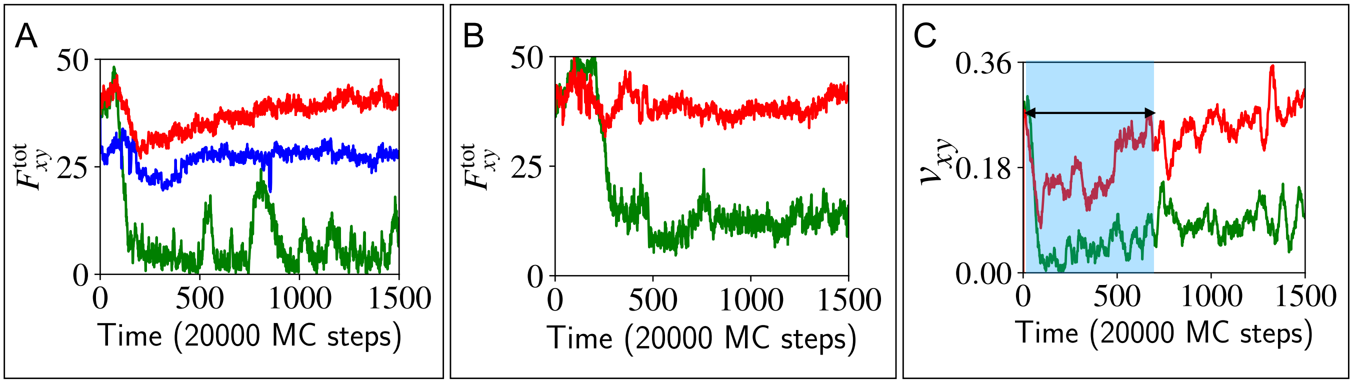


FIG. S-5. Total planar force $F_{xy}^{\rm tot}$ from the simulation is shown in (A) and (B) for rigid and soft barriers respectively. We denoted the case of $\beta = 0,\ 6,$ and 15 in units of $D/k_BT$ with the green, blue and red colours respectively. (C) The planar velocity of the vesicle during its interaction with a soft barrier for the cases $\beta = 0\ D/k_BT$ and 15 $D/k_BT$ in green and red colours respectively. It shows a similar dip in speed (denoted by a blue-shaded rectangle).

Mathematically, we can specify the wall uniquely by telling the coordinates $(w_x,\ w_y)$ through which the line passes and the angle $\xi$ with the $x$ axis as shown in Fig. S-4. The equation of this line is given by,

$$y = w_y + \tan\xi(x - w_x). \tag{S-11}$$

Let us consider a vertex move that tries to place a vertex at the coordinates $(x,\ y,\ z)$. Then, we detect the vertex is trying to penetrate the wall by the condition if $y < w_y + \tan\xi(x - w_x)$. The rigidity of a wall is characterized by a spring constant $k_w$ corresponding to the wall. Higher $k_w$ implies a more rigid wall, whereas smaller $k_w$ implies a softer wall. If the wall is completely rigid, i.e., $k_w \to \infty$ it costs infinite energy. Therefore, this vertex move is abandoned.

Besides the completely rigid walls, the wall can also be a soft one, i.e., $k_w$ is finite. Then, the energy cost for such a vertex move will be finite. Smaller $k_w$ means a softer wall. If a vertex tries to penetrate the wall (i.e., in the shaded region as shown in Fig. S-4), then the vertex feels a restoring force following Hook's law. We find the

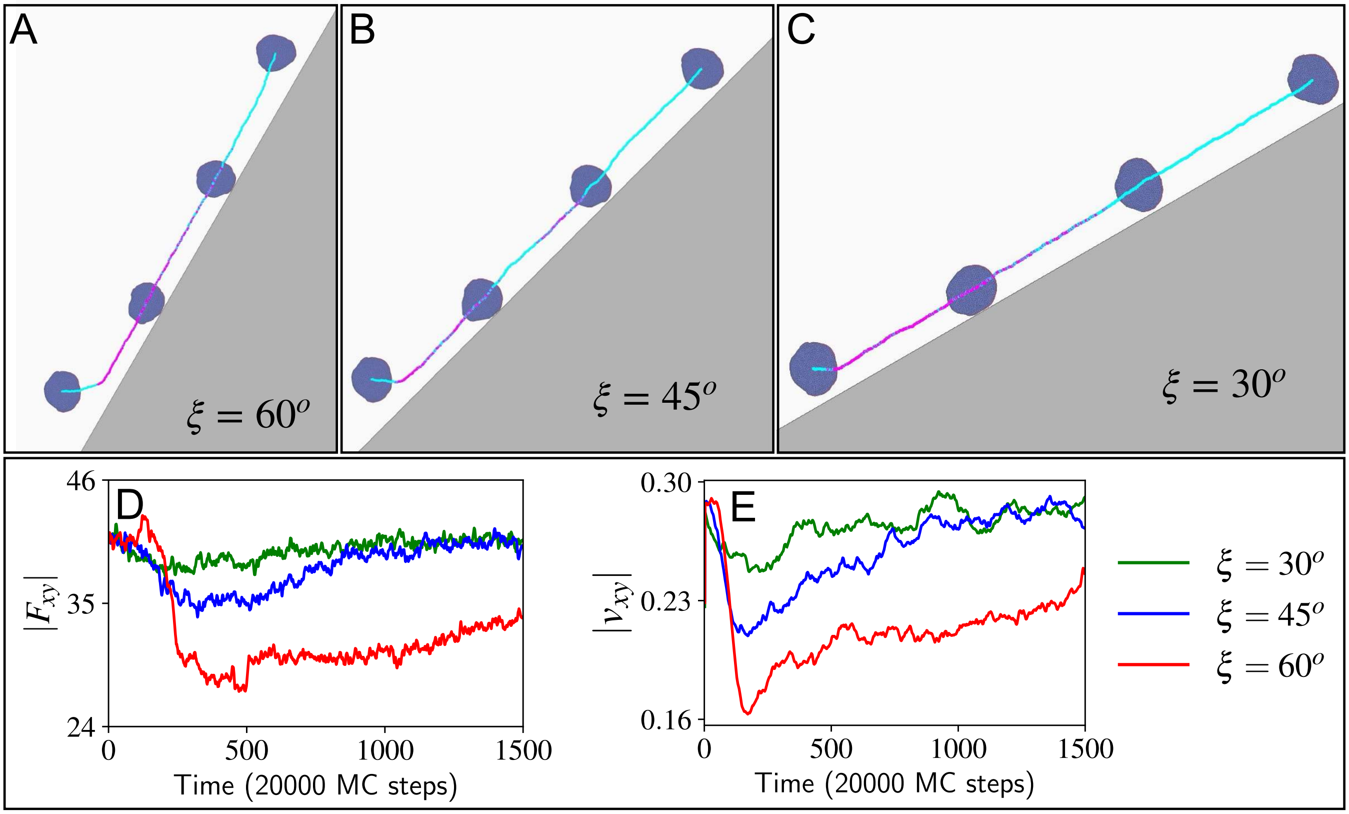


FIG. S-6. Scattering of the vesicle with slanted rigid walls ($\xi \neq 90^o$): We showed the trajectory of the vesicle scatters from the walls specified by the angles $\xi = 60^o,\ 45^o,\ 30^o$ through the point (50, 0) are shown in (a)-(c) respectively. Used parameter $E_{\rm ad} = 3k_BT,\ F = 2k_BTl_{\rm min}^{-1},\ \beta = 15\ D/k_BT$ . The magenta colour on the trajectory denotes that the vesicle touches the wall, whereas the cyan colour on the trajectory denotes that the vesicle does not touch the wall. (d)-(e) The planar force magnitude $|F_{xy}|$ and planar velocity magnitude $|v_{xy}|$ (both averaged over 5 different realizations) are shown for $\xi = 30^o,\ 45^o$, and $60^o$ with green, blue and red respectively. It shows the loss of polarity before repolarization increases as the inclination angle $\xi$ increases.

perpendicular from the vertex position (Blue dot in Fig. S-4) to the wall. Hence the soft wall will try to push back the vertex by applying a restoring force that is directed normally to the equilibrium position (red dot in Fig. S-4) of the displaced wall. The coordinate of the equilibrium position is given by, $(x - r\sin\xi, y + r\cos\xi)$, where $r = (x - w_x)\sin\xi - (y - w_y)\cos\xi$ is the perpendicular distance of the wall from the vertex when the vertex is in the grey shaded region. Therefore, the restoring force felt by the vertex is given by,

$$\begin{aligned} F_x^{\rm wall} &= -k_w r \sin\xi \\ F_y^{\rm wall} &= +k_w r \cos\xi. \end{aligned} \tag{S-12}$$

Therefore, the energy cost due to the soft wall is given by,

$$\Delta E^{\rm wall} = -F^{\rm wall}dx - F_y^{\rm wall}dy. \tag{S-13}$$

In the main text, we already discussed the results of the vesicle's head-on collision with the rigid and soft wall barriers ($\xi = 90^o$) as shown in the Fig. 5.We placed the motile vesicle in front of a slanted rigid barriers $\xi \neq 90^o$. We set the wall parameters $w_x = 50l_{\rm min},\ w_y = 0l_{\rm min}$. The typical trajectories with the angle $\xi = 30^o,\ 45^o$, and $60^o$ are shown in Fig. S-6(A-C) respectively. We set the coupling $\beta$ to 15. We see the polarity loss and repolarization through the net planar force magnitude $|F_{xy}|$ and the velocity magnitude $|v_{xy}|$ in Fig. S-6(D-E). As the angle $\xi$ increases we see the most dip in planar velocity and force increases. As the collision is more like a head-on collision it breaks the polarity more strongly. As the coupling is strong, finally they gain their polarity back.

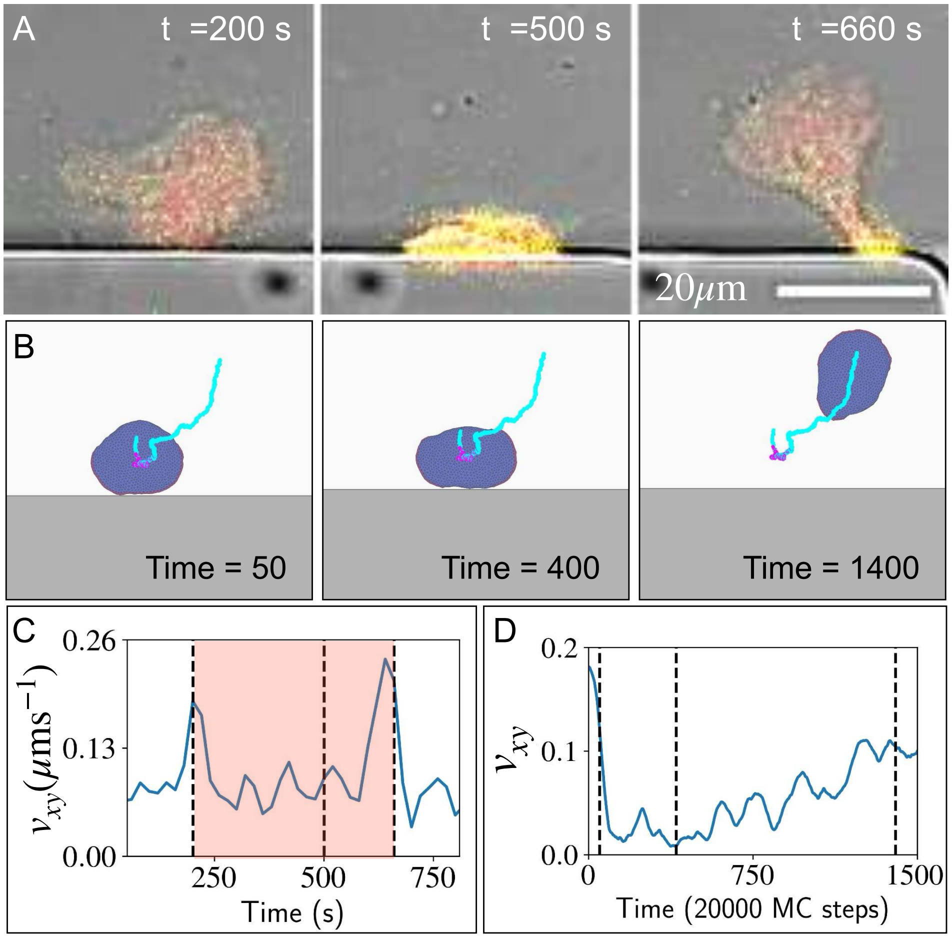


FIG. S-7. (A) Scattering of a *D. Discoideum* cell from the barrier (See Movie S-10) (B) Scattering of a polar crescent-shaped vesicle from a rigid wall ($\xi = 90^o$, $w_x = 25 l_{\rm min}$, $w_y = 0\ l_{\rm min}$) instead of hugging motion along the wall. Used parameters are $\beta = 6\ D/k_BT$, $E_{\rm ad} = 3k_BT$, $F = 2k_BTl_{\rm min}^{-1}$ (See Movie S-8). (C) The planar velocity $v_{xy}$ of the *D. Discoideum* cell in experiments. (D) The planar velocity $v_{xy}$ of our "mini-cell" in simulations.

#### G. Scattering in the non-uniform geometries

##### 1. *Hit on a triangular tip*

We placed a crescent-shaped polar vesicle in front of a triangular-tipped geometry and let it move to hit the triangular tip. To create the triangular tip geometry we allowed the vertex movement in the correct region. The triangular tip is symmetric in $y$, about the horizontal line $y = 0$. Let the sharp tipping point be located at $(x_0,\ 0)$ and the half angle of the triangular tip is $\theta$ as shown in Fig. S-8. If the position of the vertex of interest is $(x,\ y,\ z)$, then we allow the vertex to move if the new position's $x$ coordinate does not exceed a maximum value $x_{\rm max}$ for a given $y$. In addition, the maximum allowed $x$ position depends on $|y|$ as it is symmetric about $y = 0$. For a sharp triangular tip, the maximum allowed $x_{\rm max}$ is given by a linear function,

$$x_{\rm max} = |y|/\tan\theta + x_0. \tag{S-14}$$

We showed the sharp-edged triangular tip by dotted line in Fig. S-8.

To ensure the breakage of the protein aggregate smooth, we made the tip rounded near its sharp edge. If we are very close to the tipping point, we model the tipping edge as a parabolic function in such a way it should match with the linear function at $|y| = r$. Here, $r$ is the parameter with which we define the range of the rounded edge. Now, the maximum value $x_{\rm max}$ depends on the $y$ position of the vertex and it is given by,

$$x_{\rm max} = \begin{cases} \frac{|y|}{\tan\theta} + x_0, & \text{if } |y| \geq r \\ \frac{y^2/2r + r/2}{\tan\theta} + x_0, & \text{Otherwise.} \end{cases} \tag{S-15}$$

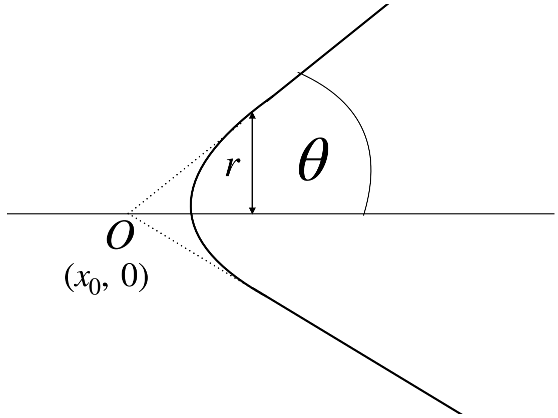


FIG. S-8. Schematic diagram of the rounded triangular tip used for the shape of the rigid barrier in Fig. 6.

We set $\theta = 45^o$ and $r = 5l_{\rm min}$ for all the simulations done in this paper. We choose a parabolic function in such a way the piece-wise derivatives match at $|y| = r$. We showed how a polar crescent-shaped vesicle loses its polarity and repolarizes in another direction while interacting with a triangular tip of a square-corner edge in Fig. 6.

#### 2. *Dumbbell-shaped confinement*

A "H" or a dumbbell-shaped confinement is implemented with rigid barriers. In doing so, we disallowed the vertex movement outside the confined region. The "H" shaped confinement is shown in the Fig. 7B. Let a vertex try to move to a position $(x,\ y,\ z)$. Then $x$ must satisfy the condition, $-Lx_2/2 \le x \le Lx_2/2$. Now, depending on $x$, we only allow those vertex moves that satisfy,

$$|x| \le Lx_2/2$$
$$|y| \le \begin{cases} Ly_1/2, & \text{if } |x| \le Lx_1/2 \\ Ly_2/2, & \text{if } Lx_1/2 \le |x| \le Lx_2/2. \end{cases} \tag{S-16}$$

The oscillation of the vesicle within such dumbbell-shaped confinement is shown in Fig. 7A. The oscillation of $x$ position and the total active force $F_x^{\rm tot}$ is shown in Fig. 7B. The breaking of the protein cluster and the reversal of the polarity are shown in Fig. 7C. Fig. 7D shows the oscillation of the curvature for the curved proteins for $x_i > x_{\rm CM}$ and $x_i < x_{\rm CM}$ in red and blue respectively. When the leading edge cluster interacts with the rigid barrier of one end it gets fattened and curvature decreases. The leading edge breaks into parts and net actin flow nearly cancels. However the protein cluster in the rear end contributes more in net actin flow and protrudes strongly, hence the curvature at the rear end increases. This mechanism leads to the oscillation of the vesicle within the dumbbell-shaped rigid confinement barrier (See Movie S-16).

### H. Dumbbell and rectangular adhesive patch

We confined the vesicle movement by placing the "mini-cell" on an adhesive patch of a dumbbell and a rectangular shape. The vesicle feels the attractive adhesion only if the vertex of the vesicle is on the adhesive portion (denoted with a grey colour) of the substrate, otherwise, the energy of adhesion for the vertex is zero (white coloured). For the dumbbell-shaped adhesive patch (See Movie S-17), we allowed the region mentioned in Eq. S-16 for adhesion. The oscillation of the vesicle on the dumbbell-shaped adhesive pattern is shown in Fig. S-9. The oscillation of the vesicle on the dumbbell-shaped adhesive substrate is shown in Fig. S-9A. In Fig. S-9B, we showed the oscillation of $x$ positions of the vesicle and the total active force along $x$ axis. Fig. S-9C shows the breaking of the protein clusters and its contribution in actin flow and finally, the repolarization in the opposite direction. The oscillation in curvature of the protein clusters belongs to right and left of the center of mass in red and blue respectively as shown in Fig. S-9D.

For the rectangular-shaped adhesive pattern, one can simply set the width $L_y = Ly_1 = Ly_2$ and length $L_x = Lx_2$. Fig. S-10 respectively. Our "mini-cell" model exhibits an oscillation (See Movie S-18) along the length of the rectangular adhesive patch as shown in Fig. S-10A. We showed the oscillation of the $x$ position (i.e., along the length of the rectangular adhesive patch) and the $x$ component of the total active force $F_x^{\rm tot}$ in Fig. S-10B. In Fig. S-10C, we showed the mean curvature of the protein sites of the vesicle for the cases $x_i > x_{\mathrm{CM}}$ and $x_i < x_{\mathrm{CM}}$ with red and blue respectively.

#### I. Cell culture and imaging

For the experimental recordings, we used non-axenic D. discoideum cells (DdB wildtype background) that are deficient in NF1, a homologue of the human RasGAP Neurofibromin [1]. They show a strongly increased tendency to switch to a highly polarized keratocyte-like mode of migration (so called fan-shaped motility) [3] that resembles the polar crescent-shaped model vesicles. The cell strain was transformed with an episomal plasmid encoding Lifeact-GFP and PHcrac-RFP (as described in [2]. Cells were cultivated in 10 cm dishes with Sørensen's buffer (14.7 mM KH2PO4, 2mM Na2HPO4, pH 6.0) supplemented with 50 µM MgCl2, 50 µM CaCl2 (Sørensen's-MC buffer) and using G418 (5 µg/ml) and hygromycin (33 µg/ml) as selection markers. Klebsiella aerogenes with an OD600 of 20 were added to the solution in 1:10 volume to a final OD600 of 2. Before imaging, cells were washed with Sørensen's-MC buffer by centrifugation at 300 x g to remove any remaining bacteria. The resulting pellet was reconstituted in Sørensen's-MC buffer and cells were left to starve for 1 hour before infusing the cell solution in the microfluidic chip using a syringe pump. Imaging was performed without flow in the microfluidic chip, using a laser scanning microscope (LSM780, Zeiss, Jena) with a 488 nm Argon laser and a 561 nm diode laser. For the experiments shown in Figures 2 and 5, acquisition was done with a 20x objective at an interval of 20 s. For the experiments shown in Figure 6, acquisition was done with a 40x oil immersion objective, at an interval of 5 s.

#### J. Fabrication of microfluidic chips

A silicon wafer coated with a 10 µm photoresist layer (SU-8 2010, Micro Resist Technology GmbH, Germany) was patterned by direct write lithography using a maskless aligner (µMLA, Heidelberg Instruments Mikrotechnik GmbH, Germany). Polydimethylsiloxane (PDMS, Sylgard 184, Dow Corning GmbH, Germany) at a ratio of 10:1 (base to curing agent) was poured into the microstructured wafer, degassed and cured for 2 h at 75°C. A PDMS block containing the microstructures was cut out and plasma bonded to a glass coverslip (#1.5, 24 × 40 mm, Menzel Glaser). The microfluidic chip was rinsed extensively with Sørensen's-MC buffer before adding the cell solution.

### S-1. MOVIE

**Breaking of the unstable crescent vesicle, stabilized by UCSP**—Crescent-shaped polar vesicle is very unstable. Simulation for four different $\beta = 0,\ 0.1,\ 0.4,$ and 20 in units of $D/k_BT$ in four columns respectively. We can see the stabilization of the vesicle as the coupling factor increases.

### S-2. MOVIE

**Breaking of the unstable crescent vesicle, higher thermal fluctuation,** $\beta = 0\ D/k_BT$—Crescent-shaped polar vesicle is very unstable. The bending rigidity of the membrane is set to $\kappa = 15k_BT$ which increases the thermal noise. It easily breaks the protein cluster and loses polarity.

### S-3. MOVIE

**Breaking due to higher thermal fluctuation, repolarize through UCSP** $\beta = 20\ D/k_BT$—Crescent-shaped polar vesicle is very unstable. The bending rigidity of the membrane is set to $\kappa = 15k_BT$ which increases the thermal noise. It breaks the protein cluster, loses polarity partially and repolarizes due to the strong UCSP coupling $\beta = 20\ D/k_BT$.

### S-4. MOVIE

**Transition from a two-arc shape to a crescent-shaped vesicle**—A two-arc shaped vesicle $F = 3k_BTl_{\text{min}}^{-1}$, $E_{\text{ad}} = 1k_BT$, can make a spontaneous transition to a crescent-shaped polar vesicle with the UCSP coupling $\beta = 10\ D/k_BT$.

### S-5. MOVIE

**Experimental: Transition from two-arc to crescent and vice versa**— Timelapse recording of a *D. discoideum* cell moving in between two PDMS barriers (rectangular shapes at the top and bottom of the field of view). The cell undergoes a transition from two-arc to crescent shape and vice versa. The time interval where the cell does not interact with the barrier and shows the transition, is presented in Fig. 2J, with t = 0 in the figure corresponding to t = 1000 s (timestamp 16:40) in the full timelapse. Green channel: LifeAct-GFP, red channel: PHcrac-RFP.

### S-6. MOVIE

**Interaction with rigid barrier, No UCSP** $\beta = 0\ D/k_BT$—A polar vesicle ($F = 2k_BTl_{\text{min}}^{-1}$, $E_{\text{ad}} = 3k_BT$) hitting a rigid barrier $\xi = 90^o$ and breaks into a non-polar two-arc shape as UCSP coupling $\beta = 0\ D/k_BT$ is absent.

### S-7. MOVIE

**Interaction with rigid barrier, UCSP** $\beta = 15\ D/k_BT$—A polar vesicle ($F = 2k_BTl_{\text{min}}^{-1}$, $E_{\text{ad}} = 3k_BT$) hitting a rigid barrier $\xi = 90^o$ and repolarize to a polar crescent shaped vesicle as UCSP coupling $\beta = 15$ is strong enough.

### S-8. MOVIE

**Scattering from the rigid barrier in large angle**—Scattering of the vesicle on the rigid barrier. We used parameters $F = 2k_BTl_{\text{min}}^{-1}$, $E_{\text{ad}} = 3k_BT$, $\beta = 6\ D/k_BT$.

### S-9. MOVIE

**Experimental: Sliding along the wall**— Timelapse recording of a *D. discoideum* cell sliding along the wall of a PDMS barrier. The corresponding trajectory and snapshots are shown in Fig. 5C. Green channel: LifeAct-GFP, red channel: PHcrac-RFP.

### S-10. MOVIE

**Experimental: Scattering in large angle**— Timelapse recording of *D. discoideum* cells moving in between two PDMS barriers (rectangular shapes at the top and bottom of the field of view). Timestamp is shown in mm:ss. Green channel: LifeAct-GFP, red channel: PHcrac-RFP.

### S-11. MOVIE

**Interaction with soft barrier, No UCSP** $\beta = 0\ D/k_BT$— A polar vesicle ($F = 2k_BTl_{\text{min}}^{-1}$, $E_{\text{ad}} = 3k_BT$) hitting a soft barrier $\xi = 90^o$. The vesicle penetrates the barrier but finally breaks into a non-polar two-arc shape as UCSP coupling $\beta = 0\ D/k_BT$ is absent.

### S-12. MOVIE

**Interaction with soft barrier, UCSP** $\beta = 15\ D/k_BT$—A polar vesicle ($F = 2k_BTl_{\min}^{-1},\ E_{\rm ad} = 3k_BT,\ \beta = 15\ D/k_BT$) hitting a soft barrier $\xi = 90^o$, is moving along the wall with a partial penetration for a longer time.

### S-13. MOVIE

**Interaction with triangular tip, No UCSP** $\beta = 0\ D/k_BT$ —A polar vesicle is placed in front of a triangular tip. The leading-edge protein cluster breaks and remains non-polar as the UCSP coupling $\beta = 0\ D/k_BT$ is absent.

### S-14. MOVIE

**Interaction with triangular tip, UCSP** $\beta = 20\ D/k_BT$—A polar vesicle is placed in front of a triangular tip. The leading-edge protein cluster breaks and loses its polarity and repolarizes itself as the UCSP coupling $\beta = 20\ D/k_BT$ is strong.

### S-15. MOVIE

**Experimental: Interaction with triangular tip**—Timelapse recording of a *D. discoideum* cell interacting with a triangular PDMS barrier. The corresponding snapshots are shown in Fig. 6D, bottom. Timestamp is shown in mm:ss. Green channel: LifeAct-GFP, red channel: PHcrac-RFP.

### S-16. MOVIE

**Oscillation on a Dumbbell patterned rigid barrier**—Oscillation of the vesicle within the dumbbell-shaped rigid barrier. We used parameters $F = 3k_BTl_{\min}^{-1},\ E_{\rm ad} = 3k_BT,\ \beta = 20\ D/k_BT,\ L_{x1} = 36l_{\min},\ L_{x2} = 78l_{\min},\ L_{y1} = 16l_{\min},\ L_{y2} = 32l_{\min}$.

### S-17. MOVIE

**Oscillation on a Dumbbell patterned adhesion**—Oscillation of the vesicle on the dumbbell-shaped adhesive pattern. We used parameters $F = 3k_BTl_{\min}^{-1},\ E_{\rm ad} = 3k_BT,\ \beta = 20\ D/k_BT,\ L_{x1} = 40l_{\min},\ L_{x2} = 80l_{\min},\ L_{y1} = 14l_{\min},\ L_{y2} = 32l_{\min}$.

### S-18. MOVIE

**Oscillation on a rectangular adhesive pattern**—Oscillation of the vesicle on the rectangular-shaped adhesive pattern. We used parameters $F = 3k_BTl_{\min}^{-1},\ E_{\rm ad} = 3k_BT,\ \beta = 20\ D/k_BT,\ L_x = 80l_{\min},\ L_y = 22l_{\min}$.

---


[1] G. Bloomfield, D. Traynor, S. P. Sander, D. M. Veltman, J. A. Pachebat, and R. R. Kay. Neurofibromin controls macropinocytosis and phagocytosis in *Dictyostelium*. eLife, 4:e04940, mar 2015.

[2] S. Flemming, F. Font, S. Alonso, and C. Beta. How cortical waves drive fission of motile cells. Proceedings of the National Academy of Sciences, 117(12):6330–6338, Mar. 2020.

[3] T. Moldenhawer, E. Moreno, D. Schindler, S. Flemming, M. Holschneider, W. Huisinga, S. Alonso, and C. Beta. Spontaneous transitions between amoeboid and keratocyte-like modes of migration. Frontiers in Cell and Developmental Biology, 10, 2022.

[4] J. E. Ron, P. Monzo, N. C. Gauthier, R. Voituriez, and N. S. Gov. One-dimensional cell motility patterns. Physical review research, 2(3):033237, 2020.

[5] R. K. Sadhu, A. Iglič, and N. S. Gov. A minimal cell model for lamellipodia-based cellular dynamics and migration. Journal of Cell Science, 136(14):jcs260744, 2023.

[6] R. K. Sadhu, S. Penič, A. Iglič, and N. S. Gov. Modelling cellular spreading and emergence of motility in the presence of curved membrane proteins and active cytoskeleton forces. The European Physical Journal Plus, 136(5):495, 2021.

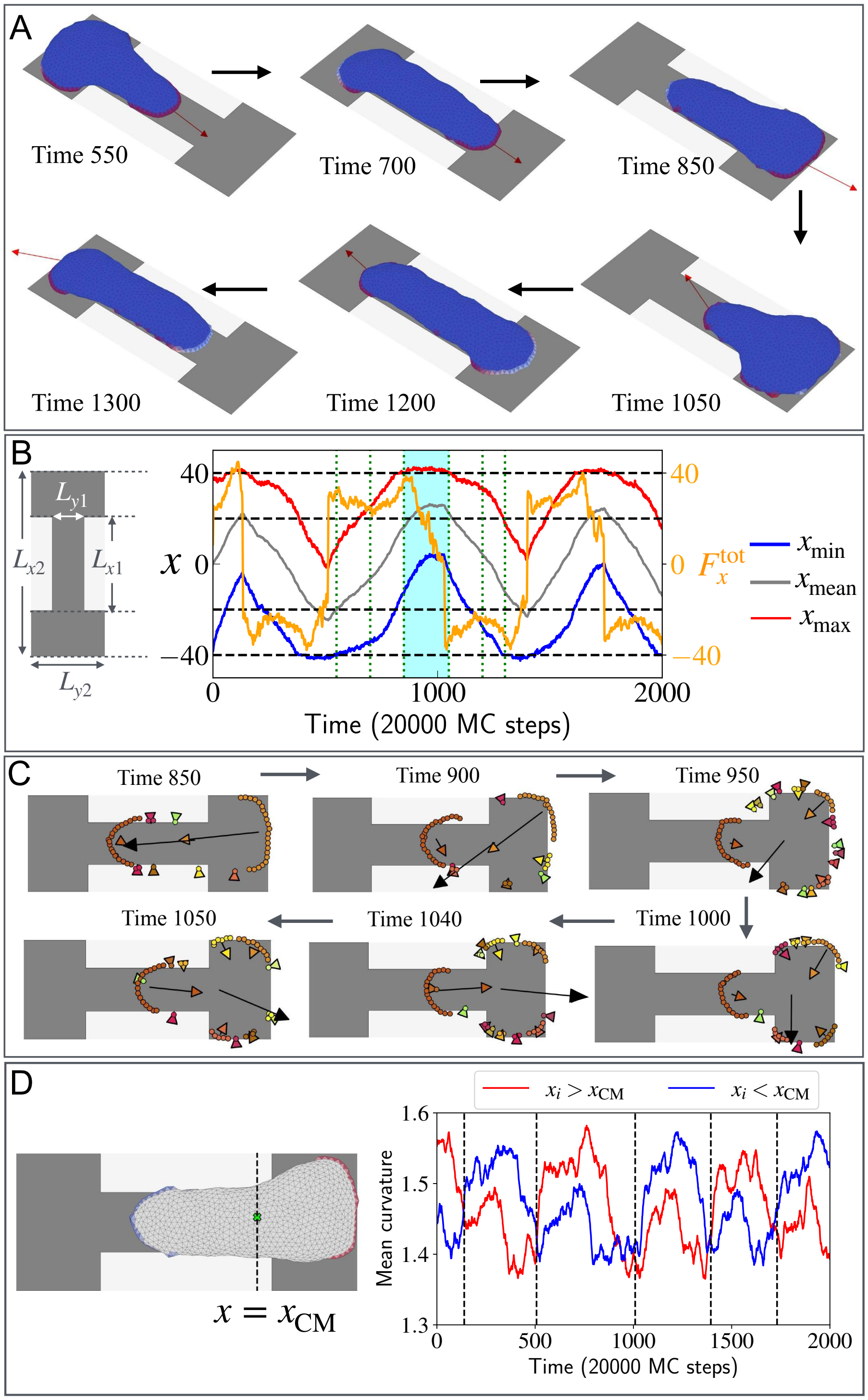


FIG. S-9. Spontaneous oscillations of a vesicle confined on a dumbbell-shaped adhesion patch. (A) Snapshots of the vesicle at different times, during a complete oscillation between the two chambers. The dynamics of the total planar active force $\boldsymbol{F}_{xy}^{\text{tot}}$ is indicated by the red arrow. We use the adhesion strength $E_{ad} = 3k_BT$, active force parameter $F = 3k_BTl_{\text{min}}^{-1}$, and the coupling parameter $\beta = 20\ D/k_BT$. (B) The oscillation of the vesicle's $x_{\text{min}}$, $x_{\text{mean}}$, and $x_{\text{max}}$ of all the vertices (blue, grey, and red solid lines). The dimensions of the dumbbell shaped-confinement: $L_{x1} = 40$, $L_{x2} = 80$, $L_{y1} = 14$, and $L_{y2} = 32$ in the units of $l_{\text{min}}$. The time evolution of the $x$ component of the total force $F_x^{\text{tot}}$ is shown, with the scale given on the right axis in orange. (C) Different protein clusters are shown in different colours on a $x - y$ plane from the top view during the polarity flip. We indicate the actin flow contribution from each cluster using a correspondingly coloured arrow. The total actin flow is indicated using a black arrow at the centre of mass of the vesicle. (D) A schematic diagram of the vesicle indicates that the CMC are coloured red for $x_i > x_{\text{CM}}$, and blue for $x_i < x_{\text{CM}}$. On the right, the dynamics of the mean curvature for the red and blue CMC is shown. The black dashed lines denote the reversals of the total force along the $x$ direction (B).

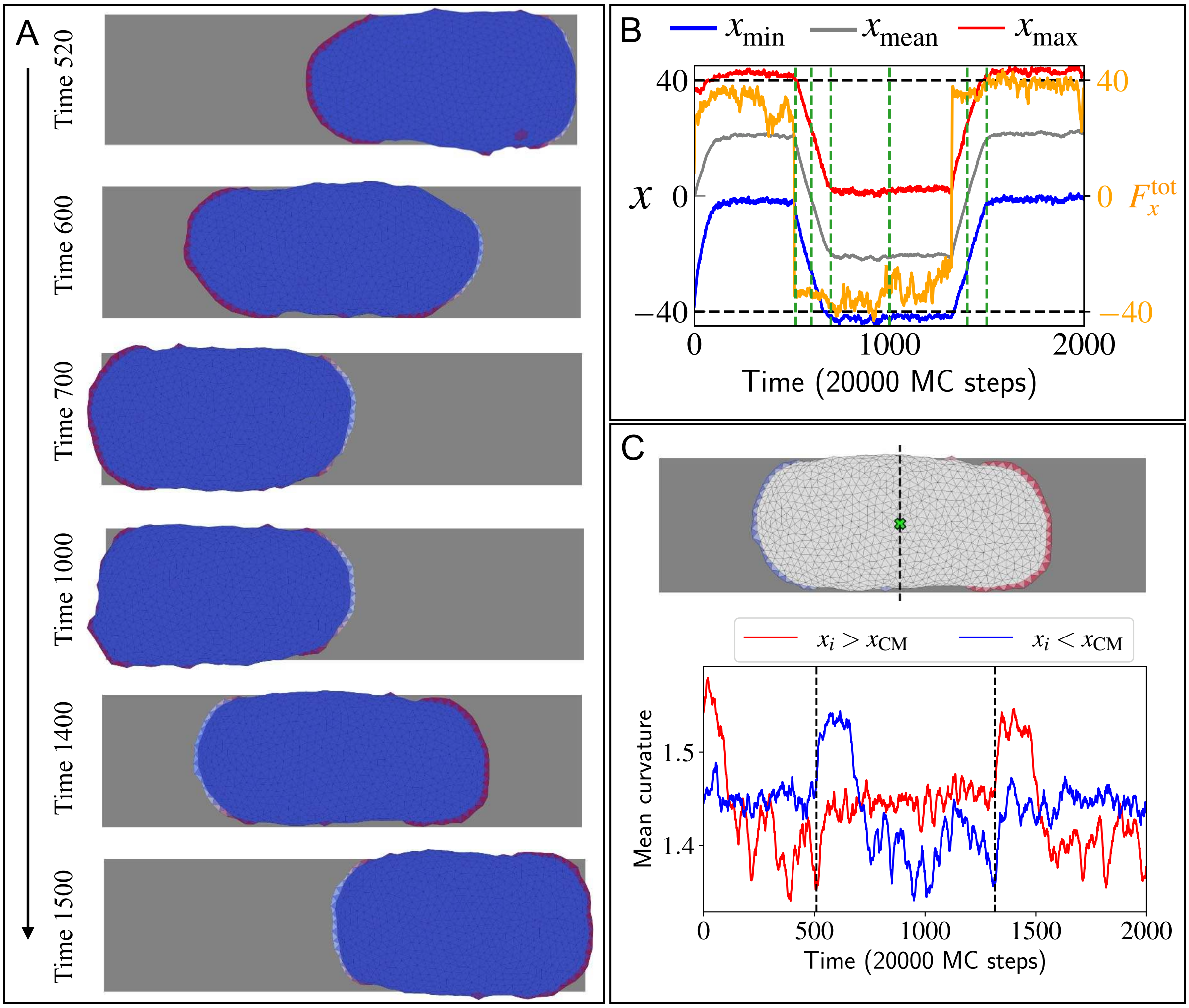


FIG. S-10. (A) The snapshots of the vesicle within the dumbbell-shaped confinement at time 520, 600, 700, 1000, 1400, and 1500 in the units of 20000 MC Steps during a complete oscillation on the rectangular adhesive substrate. The length and the width of the rectangle are $80l_{\mathrm{min}}$ and $22l_{\mathrm{min}}$ respectively. (B) The oscillation in $x$ coordinates of the vesicle over time. The $x_{\mathrm{min}}$, $x_{\mathrm{mean}}$, and $x_{\mathrm{max}}$ of all the vertices of the vesicle are shown in blue, grey, and red solid lines. The time evolution of the $x$ component of the total force $F_x^{\mathrm{tot}}$ is shown in the dual scale in orange. Green vertical dashed lines denote the time of the snapshots in (A). (C) On the top, a schematic diagram of the vesicle where the curved proteins are coloured with red if it's $x$ coordinate $x_i > x_{\mathrm{CM}}$, and with blue if $x_i < x_{\mathrm{CM}}$. The centre of mass is denoted by a lime-coloured marker. On the bottom, the mean curvature for the curved proteins for $x_i > x_{\mathrm{CM}}$ and $x_i < x_{\mathrm{CM}}$ in red and blue respectively. The black dashed lines denote the reversal of force along the $x$ direction.
